# MethReg: estimating the regulatory potential of DNA methylation in gene transcription

**DOI:** 10.1101/2021.02.18.431696

**Authors:** Tiago C. Silva, Juan I. Young, Eden R. Martin, Xi Chen, Lily Wang

## Abstract

Epigenome-wide association studies (EWAS) often detect a large number of differentially methylated sites or regions, many are located in distal regulatory regions. To further prioritize these significant sites, there is a critical need to better understand the functional impact of CpG methylation. Recent studies demonstrated CpG methylation-dependent transcriptional regulation is a widespread phenomenon. Here we present MethReg, an R/Bioconductor package that analyzes matched DNA-methylation and gene-expression data, along with external transcription factor (TF) binding information, to evaluate, prioritize, and annotate CpG sites with high regulatory potential. By simultaneous modeling three key elements that contribute to gene transcription (CpG methylation, target gene expression and TF activity), MethReg identifies TF-target gene associations that are present only in a subset of samples with high (or low) methylation levels at the CpG that influences TF activities, which can be missed in analyses that use all samples. Using real colorectal cancer and Alzheimer’s disease datasets, we show MethReg significantly enhances our understanding of the regulatory roles of DNA methylation in complex diseases.

## Background

DNA methylation is a widely studied epigenetic modification. Recent epigenome-wide association studies (EWAS) have identified numerous alterations in DNA methylation (DNAm) levels that are involved in many diseases such as various cancers [1–5] and neurodegenerative diseases [6–8]. Compared to genome-wide association studies (GWAS) of genetic variants, EWAS often detect a larger number of significant differences, often thousands of differentially methylated CpG sites (DMS), which are significantly associated with a disease or phenotype. Many of these DMS are located far from genes, complicating the interpretation of their functionality [9, 10]. Therefore, there is a critical need to better understand the functional impact of these CpG methylations and to further prioritize the significant methylation changes.

Transcription factors (TFs) are proteins that bind DNA and facilitate the transcription of DNA into RNA. A number of recent studies have observed that the binding of TFs onto DNA can be affected by DNA methylation, and in turn, DNA methylation can also be altered by proteins associated with TFs [11–15]. Using methylation-sensitive SELEX (systematic evolution of ligands by exponential enrichment), Yin et al. (2017) [16] classified 519 TFs into several categories: TFs whose binding strength increased, decreased, or was not affected by DNA methylation, as well as those not containing CpGs in their binding motifs.

Although a number of integrative analyses strategies [17–23] have been proposed to help assess the functional role of the DNA methylation changes in gene regulation, these methods typically integrate DNA methylation data with either gene expression [17–20] or TF binding data [21–23], but rarely both. For example, MethylMix [17] identifies CpGs that are predictive of transcription and then classifies the CpGs into different methylation states. Similarly, COHCAP [18] identifies a subset of CpGs within CpG islands that are most likely to regulate downstream gene expression. These methods which test the association between DNA methylation and target gene expression can be further improved by additionally incorporating information on TF activities.

To determine whether TF regulatory activity is enhanced or reduced by significant CpGs in EWAS, the gold standard would be to perform ChIP-Seq experiments for all candidate TFs with binding sites close to the CpGs in parallel with bisulfite sequencing of DNA methylation changes and transcriptome assessment.

However, performing ChIP-Seq experiments on primary tissues is technically challenging because of the limited number of cells in some samples, the large number of TF to be tested, and the lack of availability for specific antibodies. Therefore, in practice, computational approaches are often used to prioritize disease relevant TFs in DNA methylation studies. For example, LOLA [21] (Locus Overlap Analysis) performs enrichment analysis to identify regulatory elements such as TFs with binding sites enriched in candidate genomic regions (e.g., DMRs). Alternatively, Goldmine [22] annotates individual CpGs by TFs with binding sites that overlap with the CpG. However, because the binding motifs for TFs are often non-specific for different members of a TF family, there are often many TFs with binding sites that overlap with a given CpG. Moreover, TF binding can also occur without affecting the transcription of any gene [24, 25]. Therefore, methods which analyze DNA methylation and TF binding sites (TFBS) data would also be greatly enhanced by additionally modeling target gene expression.

To fill this critical gap in analytical methods and software for annotating and prioritizing DNA methylation changes identified in EWAS, here we present MethReg, an R/Bioconductor package that performs integrative modeling of three key components (DNA methylation, gene expression levels, TF) in gene regulation, to provide a more comprehensive functional assessment of CpG methylation in gene regulation. In particular, MethReg leverages information from external databases on TFBS, ChIP-seq experiments and TF-target interactions, performs both promoter and distal (enhancer) analyses, implements rigorous robust regression models, and can fully adjust for potential confounding effects such as copy number, age, and sex that are important in DNA methylation analysis. MethReg can be used either to evaluate the regulatory potentials of candidate regions identified in EWAS (in supervised analysis mode) or to search for methylation dependent TF regulatory processes in the entire genome (in unsupervised analysis mode).

Using simulated datasets, we showed that by simultaneous modeling of three key elements (DNA methylation, target gene, and TF), MethReg significantly improves prioritization for true positive DNA methylation changes with regulatory roles in gene transcription when compared to models that include only two key elements. In addition, we also analyzed the TCGA colorectal datasets and the ROSMAP Alzheimer’s dataset to show that MethReg was able to recover known biology, as well as nominate novel biologically meaningful DNA methylation-TF-target associations that influence transcription.

## Results

### An overview of the MethReg software

To systematically search for CpG methylation with significant regulatory effects on gene expression by influencing TF activities, we developed MethReg. Figure 1 illustrates the workflow for MethReg. Given matched (methylation arrays) DNA methylation data and (RNA-seq) gene expression data, MethReg additionally incorporates TF binding information from ReMap2020 [26] or the JASPAR2020 [27] database, and optionally additional TF-target gene interaction databases (Supplementary Table 1), to perform both promoter and distal (enhancer) analysis. In the unsupervised mode, MethReg analyzes all CpGs on the Illumina arrays. In supervised mode, MethReg analyzes and prioritizes differentially methylated CpGs (DMS) identified in EWAS. There are three main steps: (1) create a dataset with triplets of CpGs, TFs which bind near the CpGs, and putative target genes; (2) for each CpG-TF-target gene triplet, apply integrative statistical models to DNA methylation, target gene expression, and TF activity values; and (3) visualize and interpret results from statistical models to estimate the impact of DNA methylation on TF activity (synergistic effect of CpG methylation and TF on target gene), as well as to annotate the roles of TF and CpG methylation in regulating target gene expression. The results from the statistical models will allow us to identify a list of CpGs that work synergistically with TFs to influence target gene expression. Next, we discuss each of these analytical steps in detail. We will describe the analysis of TFs, but the methods and software tool are, in principle, also applicable to other types of chromatin proteins that crosstalk with DNA methylation. MethReg is an open-source R/Bioconductor package, available at https://bioconductor.org/packages/MethReg/.

**Figure 1.**
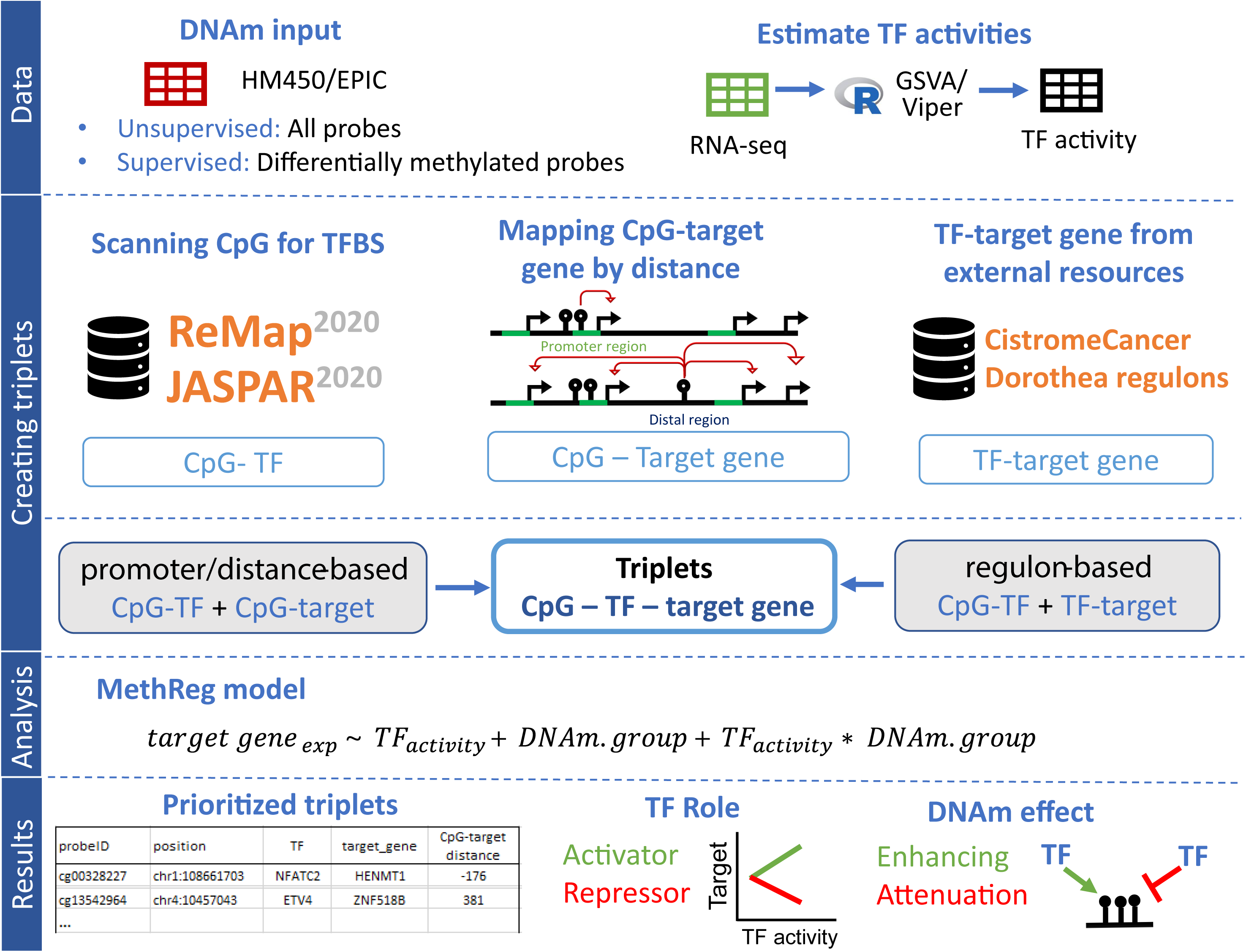
Analysis workflow of MethReg.

### Creating CpG-TF-target gene triplet dataset

MethReg first links CpGs to TFs with binding sites within a window of user-specified distance (e.g., ± 250 bp) using information from ReMap2020 [26] or JASPER2020 [27] database. The JASPER2020 [27] database includes curated transcription factor binding models, among which 637 are associated with human TFs with known DNA-binding profiles [28]. Similarly, the human atlas of the ReMap2020 database [25] contains regulatory regions for 1135 transcriptional regulators obtained using genome-wide DNA-binding experiments such as ChIP-seq. Next, in *promoter analysis*, CpGs located in promoter regions, defined as ± 2 kb regions around the transcription start sites (TSS), are linked to target genes with promoters that overlap with the CpG. On the other hand, CpGs in the distal regions (i.e., > 2kb from TSS) are linked to a specific number of genes (e.g., 5 genes) upstream or downstream and within 1M bp of the CpG. The CpG-TF pairs are then combined with CpG-target pairs to create triplets of CpG-TF-target genes.

Alternatively, CpGs can also be linked to genes within 1M bp using *regulon-based analysis*. A TF regulon consists of all the transcriptional targets of the TF. MethReg obtains TF-target pairs from curated external regulon databases [29–31] (Supplementary Table 1). Combining the CpG-TF pairs with TF-target gene pairs, we then obtain a triplet dataset where each row contains identifiers for a CpG, a TF, and the target gene.

### Estimating regulatory effects of CpG methylation and TFs on target gene expression

Given a CpG-TF-target gene triplet, we then query the matched DNA methylation and gene expression datasets to obtain DNA methylation, target gene, and TF gene expression (or activity) values and fit the following statistical model to data:

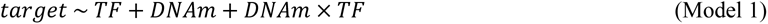

where *target* is log_2_ transformed target gene expression values, *TF* is log_2_ transformed TF gene expression values or estimated TF activity scores (see details in section “Modeling TF protein activities” below), and *DNAm* is DNA methylation beta-values.

Note that Model 1 partitions the effects of DNA methylation and TF on target gene expression into three categories: the direct effect of TF (modeled by term *TF*), the direct effect of DNA methylation (modeled by term *DNAm*), and the synergistic effects of TF and DNA methylation (i.e., how the effect of TF on target gene expression is modified by DNA methylation; modeled by *DNAm × TF* interaction term). For accurate statistical modeling, MethReg implements Model 1 by fitting a robust linear model which gives outlier gene expression values reduced weights [32]. Note that a key feature of Model 1 is that it provides more comprehensive modeling of gene regulation by incorporating the three components (TF activity, DNA methylation, target gene expression) simultaneously. In addition to Model 1 described above, which included *DNAm* as a continuous variable, we also considered another model that modeled methylation values as a binary variable. We also propose:

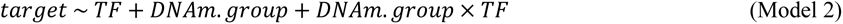

where *DNAm.group* is high or low. That is, for a given CpG, the samples with the highest DNA methylation levels (top 25%) have *DNAm.group* = “high”, and samples with lowest DNAm levels (bottom 25%) have *DNAm.group* = “low”. In this model, only samples with DNA methylation values in the first and last quartiles are considered. Note also that statistically, the *DNAm.group* x *TF* effect is estimated by comparing the magnitude of TF-target gene association in high methylation group versus the magnitude of the TF-target gene association in low methylation group.

### Modeling TF protein activities

Given that TF gene expression might not accurately reflect TF protein activities, which involve additional complex processes (e.g. post-translational modifications, protein-protein/ligand interactions, and localization changes), MethReg implements an additional option to model TF activities via the VIPER (virtual Inference by enriched regulon analysis) [33] or GSVA [34] methods, so that the TF effects in Model 1 and Model 2 described above can also be computed by replacing TF gene expression levels with estimated TF activities. Briefly, given RNA-seq data, these methods estimate the activity of a TF by performing a rank-based gene set enrichment analysis of its target genes (i.e., its regulon). MethReg can work with different regulon databases (Supplementary Table 1), such as those described Garcia-Alonso et al. (2019) [29], which were collected from four resources: manually curated databases, ChIP-seq binding experimental data, prediction of TF binding motifs based on gene promoter sequences, or computational regulatory network analysis. The Genotype-Tissue Expression (GTEx) tissue-consensus regulons included 1,077,121 TF–target candidate regulatory interactions between 1,402 TFs and 26,984 targets, and the pan-cancer regulons included 636,753 TF–target candidate regulatory interactions between 1,412 TFs and 26,939 targets. Garcia-Alonso et al. (2019) annotated each TF-target interaction with a five-level confidence score, with “A” indicating most reliable, supported by multiple lines of evidence, and “E” indicating least confidence, supported only by computational predictions. Benchmark experiments using three separate datasets showed the GTEx tissue-consensus regulons performed similarly to tissue-specific regulons computed from GTEx data of specific tissue type. Notably, MethReg also provides options for users to input alternative TF regulon databases (Supplementary Table 1) and TF activities computed using alternative software such as Lisa [35].

### Stage-wise method for controlling false discovery rate

MethReg implements two alternative methods for controlling false discovery rates, using the conventional approach [36] or a stage-wise approach [37]. To help improve power in high-throughput experiments where multiple hypotheses are tested for each gene, van de Berge et al. (2017) [37] proposed a stage-wise approach in the context of gene splicing analysis. First, in the screening step, a global test is applied to each gene to test the null hypothesis that there is a differential change in any of the transcripts within the gene. Second, in the confirmation step, for the genes selected in the screening step, individual transcripts are then tested while controlling family-wise error rate (FWER). By aggregating effects from individual transcripts within a gene in the screening step, the stage-wise procedure was shown to have superior power compared with the conventional approach that tests all individual transcripts in one-step.

In Model 1 and Model 2 described above, the synergistic activity of DNA methylation and TF is estimated by the interaction term *DNAm × TF*. Because the standard error of interaction effects is typically much larger than those for main effects, the conventional approach for controlling false discovery rate often results in low power for discovering interaction effects [37]. To this end, MethReg additionally implements the stage-wise procedure for testing interactions by first aggregating all CpG-TF-target triplets associated with the same CpG as a group. In the screening step, MethReg tests the null hypothesis that any of the individual triplets mapped to a CpG has a significant *DNAm × TF* effect. In the confirmation step, MethReg tests all the triplets associated with significant CpGs selected in the screening step while controlling FWER as described in Van den Berge et al. (2017) [37].

### Visualizing and annotating roles of CpG and TF in gene transcription

To visualize how DNA methylation and TFs work together to influence gene expression, MethReg generates a suite of figures. Figure 2 shows an example output figure of Model 2 applied to the TCGA colorectal cancer dataset (Online Methods). In the first row are figures for assessing direct pairwise TF-target and DNA methylation-target associations. In the second row are figures for assessing TF-target gene expression, stratified by high or low DNA methylation levels. In the third row, we stratify by TF expression (or activity score) instead and plot gene expression against DNA methylation values, stratified by high or low TF expression (or activity) levels.

**Figure 2.**
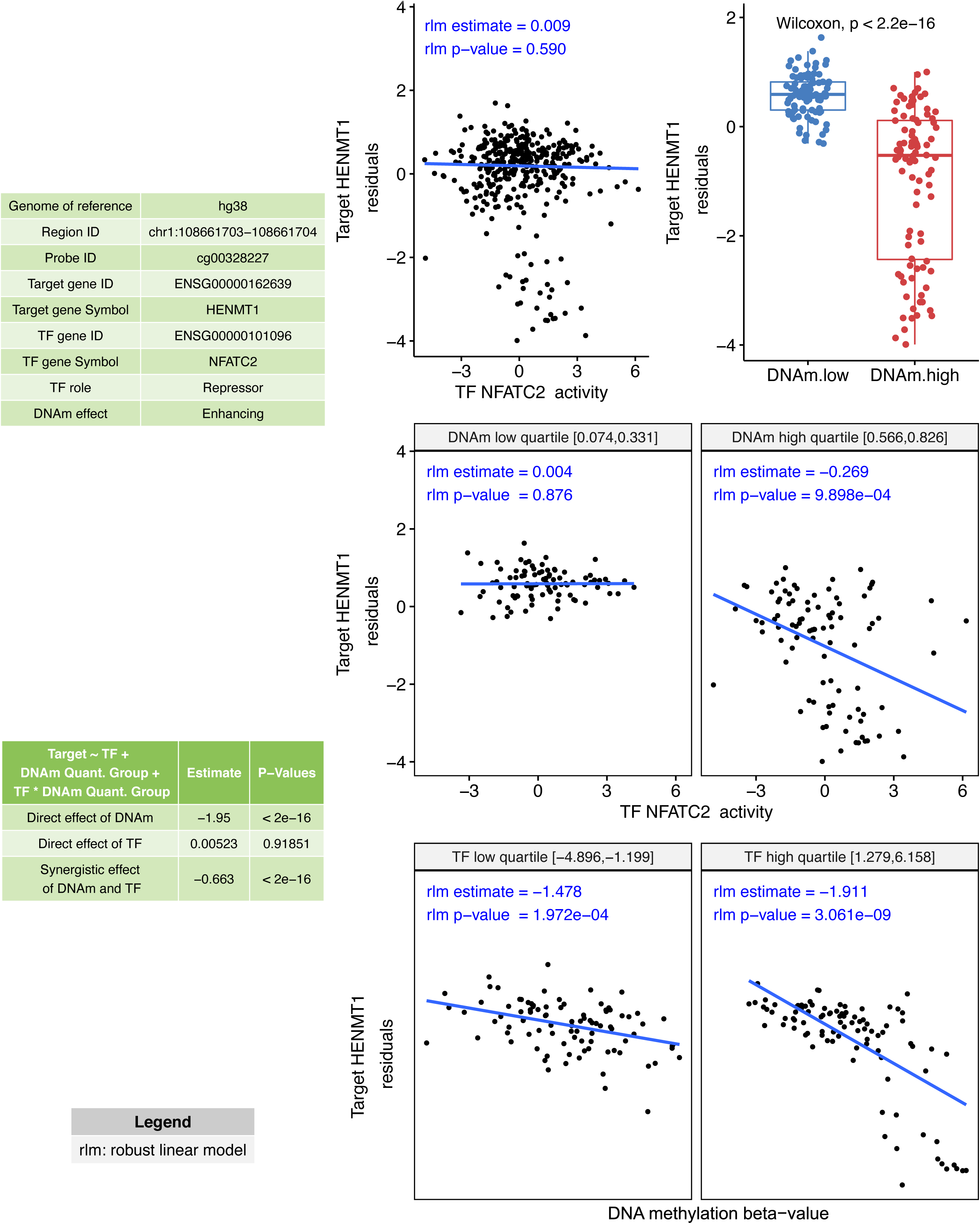
An example output from MethReg unsupervised analysis of TCGA COAD-READ smaples, in which CpG methylation enhances TF activity.

Note in Figure 2, without stratifying by DNA methylation, the overall TF-target effect (robust linear model P-value = 0.590) is not significant. In contrast, TF-target association is highly significant in samples with high methylation levels (robust linear model P-value = 0.001). Therefore, methylation at cg00328227 plays an important role in regulating TF activity in this case. This example also demonstrates that by additionally modeling DNA methylation, we can nominate TF-target associations that might have been missed otherwise.

Figure 3 shows the different biological scenarios where methylation and TF influences target gene expression synergistically. A TF repressor decreases transcription while a TF activator increases it, and the presence of methylation can either enhance or attenuate the binding affinity of the TF. For each triplet, MethReg annotates the role of TF on the target gene (repressor, activator, or dual) and how DNA methylation influences the TF (enhancing, attenuating, or invert).

**Figure 3.**
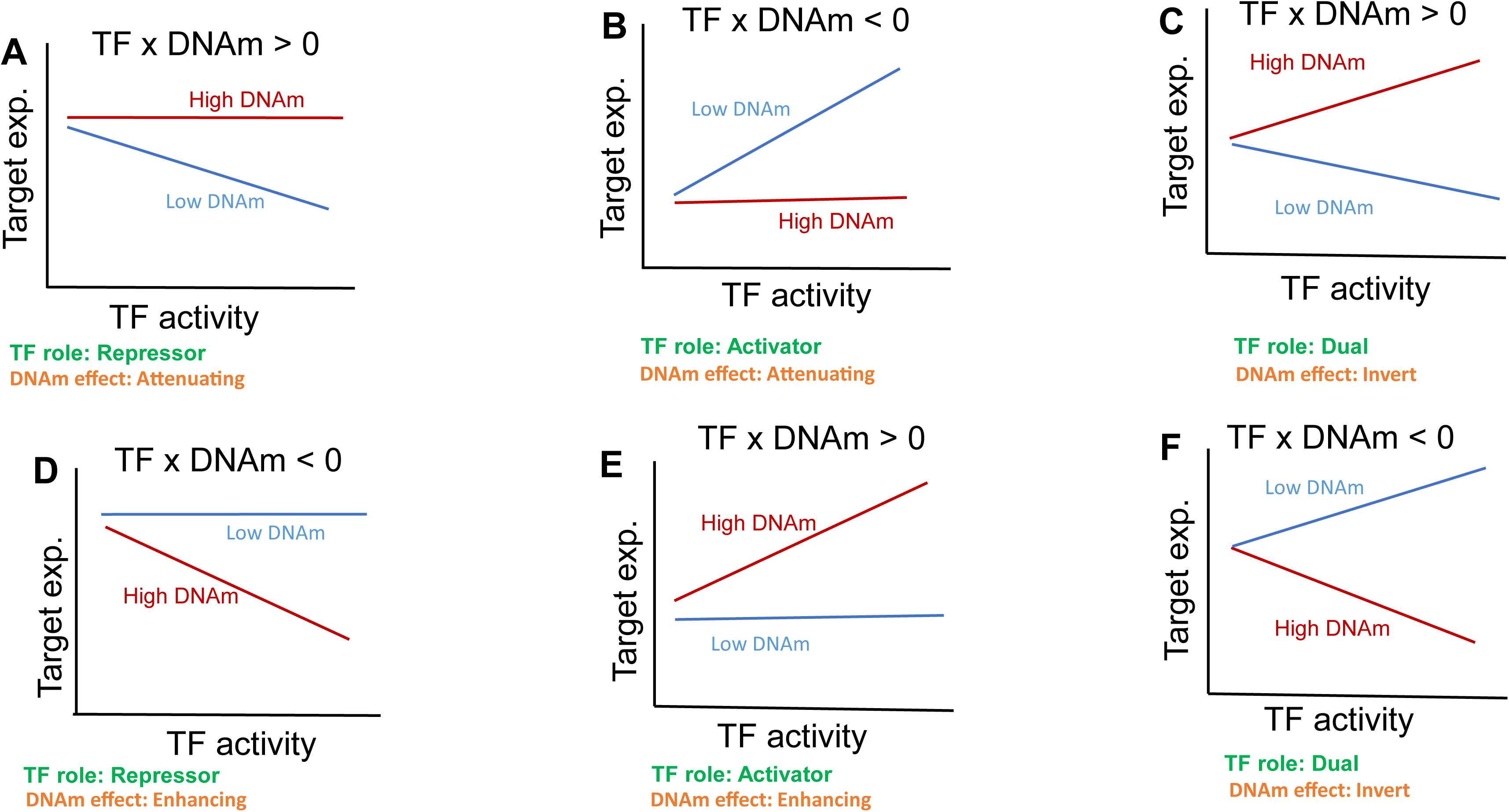
Scenarios modeled by MethReg.

### Simulation analysis

We conducted a simulation study to compare the performance of 7 different approaches, including Model 1 & Model 2 as well as the conventional approach of directly correlating TF and target gene expressions, to identify TF-target associations where TF activities are modulated by CpG methylation (Supplementary Figure 1). We considered the scenario where CpG methylation affects TF binding affinity, so that TF only affects target gene expression when methylation level is low. To assess the statistical properties of these different methods, we estimated and compared type I error rate, power, and area under receiver operating characteristics curves (AUC) for each method.

More specifically, we simulated datasets for which target gene expression levels were dependent on TF expressions only in samples with low DNA methylation levels. We used 38 samples of TCGA COAD matched RNA-seq and DNA methylation data on chromosome 21 included in the MethReg R package as our input dataset, from which we randomly sampled TF gene expression and methylation expression levels. For each simulated triplet dataset, we randomly sampled one gene from RNA-seq dataset and one CpG site from methylation dataset to be our TF expression and DNA methylation expression. Next, target gene expression levels were simulated from negative binomial distributions as follows: the estimated median mean and variance over all genes in our input dataset were *μ*_0_ = 10.59 and *σ*^2^ = 16.90, respectively. Therefore, for target gene expressions, we assumed a negative binomial distribution with parameters *μ*_0_ = 10.59 and *k*_0_ = *μ*_0_^2^/(*σ*^2^ − *μ*_0_) = 17.78 for all samples except those with lowest DNA methylation levels in the first quartile. For the samples with low methylation levels in the first quartile, we generated target gene expression levels from negative binomial distribution (*μ* = *μ*_0_ + *β* × TF gene expression, *k* = *k*_0_), where *β* = (0, 1, 2, … , 9) indicates different strengths of associations between TF and target gene expressions, corresponding to 10 different simulation scenarios. Therefore, by design of this simulation experiment, target gene expressions were associated with TF only when methylation levels were low. For each simulation scenario, we repeated this process 1,000 times to generate 1,000 triplets of TF, DNA methylation, and target gene expression levels.

Note that when *β* = 0, target gene expressions were generated randomly from negative binomial distribution and did not depend on TF gene expression in the samples, so the 1,000 triplet datasets simulated under this simulation scenario (null triplets) allowed us to estimate type I error rates of different models. We compared sensitivity and specificities for identifying TF-target associations based on p-values from 7 different approaches (Supplementary Figure 1):

**a)** lm.cont: p-value for *DNAm × TF* term in linear model implementation of Model 1.
**b)** lm.binary: p-value for *DNAm.group × TF* term in linear model implementation of Model 2.
**c)** rlm.cont: p-value for *DNAm × TF* term in robust linear model implementation of Model 1.
**d)** rlm.binary: p-value for *DNAm.group × TF* term in robust linear model implementation of Model 2.
**e)** rlm.binary.en: p-value for *DNAm.group × TF* term in robust linear model implementation of Model 2, estimated from empirical null distribution [38] (see details in Methods).
**f)** lm.main.tf: p-value for *TF* term in main effect model *target gene expression ∼ TF*.
**g)** spearman.corr.tf: p-value for spearman correlation between target gene expression and TF.

Note that the last two methods lm.main.tf and spearman.corr.tf evaluate the conventional approach of directly correlating target gene expression with TF expression, without considering DNA methylation (Supplementary Figure 1). Also, in method e) rlm.binary.en, instead of the conventional approach which computes P-values by comparing test statistic for *DNAm.group × TF* to *t-*distribution, this method estimates P-values for *DNAm.group × TF* effect using empirical null distribution [38], which is a normal distribution with empirically estimated mean 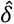 and standard deviation 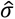. Efron (2010) [38] showed that in large-scale simultaneous testing situations (e.g., when many triplets are tested in an analysis), serious defects in the theoretical null distribution may become obvious, while empirical Bayes methods can provide much more realistic null distributions.

The results showed that all methods had type I error rates close to 5%. In particular, the methods lm.main.tf, rlm.binary.en, and spearman.corr.tf had type I error rates below 5% (Supplementary Figure 2). Among the methods, rlm.binary and rlm.binary.en had similar and the highest power across all simulation scenarios, while the main effect models that test association between target gene expression and TF expression using linear model (lm.main.tf) or Spearman correlation (spearman.corr.tf) lacked power (Supplementary Figure 3).

Given the known status of CpG methylation’s role in association with target gene expression (i.e. true negative when β = 0 and true positive when β > 0), we next computed the area under the ROC curve (AUC) for each method. The receiver operating characteristic (ROC) curves show a trade-off between sensitivity and specificity as the significance cutoff is varied. AUC assesses the overall discriminative ability of the methods to determine whether a given methylation CpG is driving target gene expression over all possible cutoffs. The best performing methods with highest AUCs are methods rlm.binary (0.894), rlm.binary.en (0.882), and lm.binary (0.838), followed by p-values from models with continuous methylation levels, lm.cont (0.821) and rlm.cont (0.815) (Figure 4). Again, the main effect models lm.main.tf and spearman.corr.tf lacked sensitivity (i.e., power) therefore had the lowest AUCs at 0.631 and 0.582, respectively.

**Figure 4.**
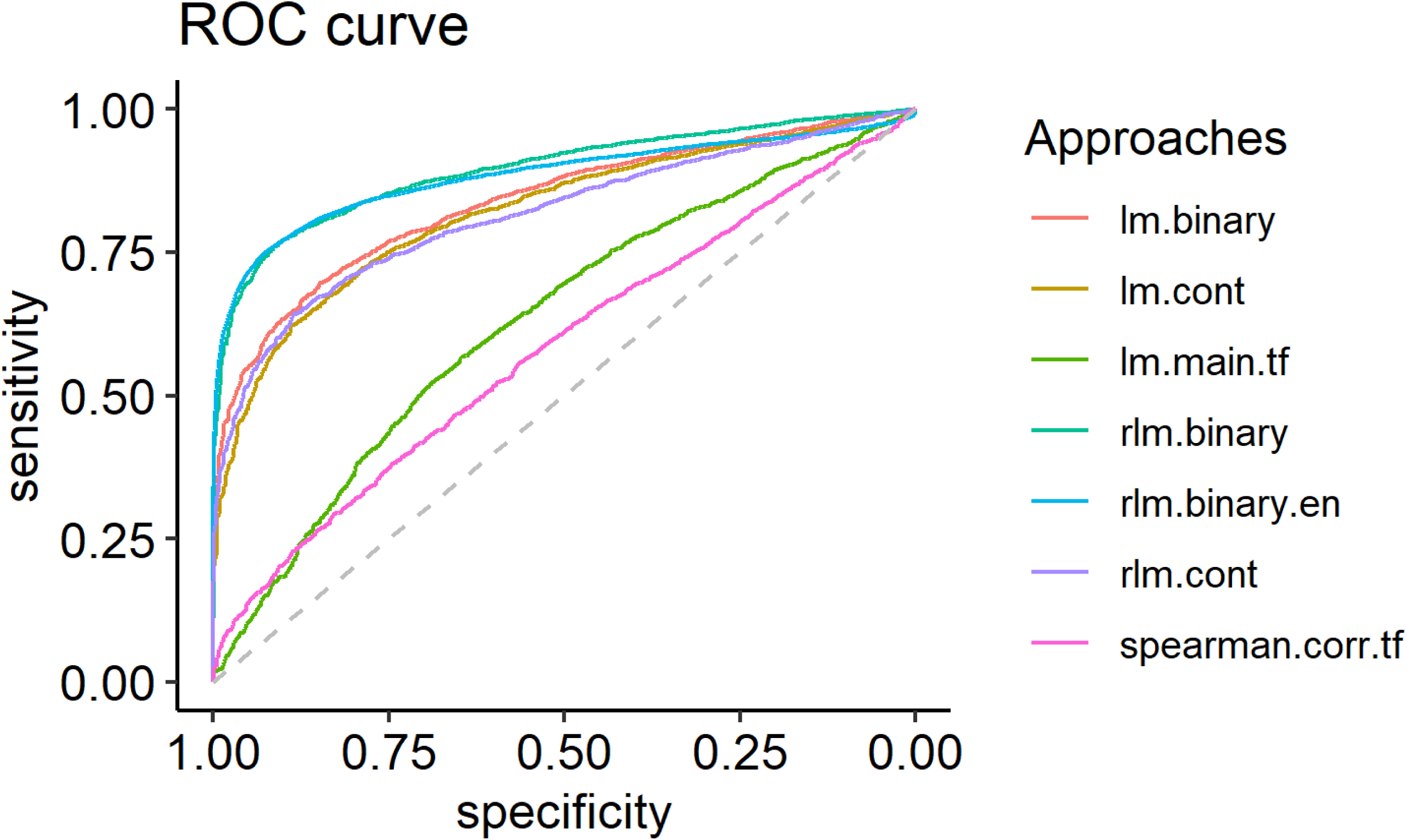
Receiver Operating Characteristics curves for different methods compared in simulation study. lm.binary = linear model implementation of Model 2; lm.cont = linear model implementation of Model 1; lm.main.tf = linear model gene expression ∼ TF; rlm.binary = robust linear model implementation of Model 2; rlm.binary.en = same as rlm.binary model, expect P-values were estimated using empirical null distribution; rlm.cont = robust linear model implementation of Model 1; spearman.corr.tf = Spearman correlation between gene expression and TF expression; Model 1: gene expression ∼ TF + DNAm+TF*DNAm; Model 2: gene expression ∼T F + DNAm.group+ TF*DNAm.group

These simulation study results showed that the conventional approach of directly correlating gene expression with TF expression (lm.main.tf and spearman.corr.tf) lacked power for detecting TF activities on target gene expression when they are also modulated by DNA methylation. This is probably because without stratifying on DNA methylation levels, the main effect models detect TF-target gene associations averaged over all samples with various methylation levels. In contrast, the interaction terms in Model 1 & 2 compare the magnitude of TF-target gene association in low methylation samples versus the magnitude of the association in high methylation samples.

The results also indicated that the models that use a binary variable (low, high) to model methylation levels (rlm.binary, rlm.binary.en and lm.binary methods) performed best, probably because these models can reduce noise in data and thus can improve power. Among binary models, the robust linear model rlm.binary and rlm.binary.en performed similarly and better than regular linear model lm.binary. Thus, we selected the rlm.binary model for our subsequent analyses.

### Case Study: analyses of TCGA COAD-READ datasets

#### An unsupervised MethReg analysis

Colorectal cancer (CRC) is the third most commonly diagnosed cancer and the second leading cause of cancer death in the United States [39]. CRC, like many other cancers, is characterized by global hypo-methylation leading to oncogene activation, chromosomal instability, and locus-specific hyper-methylation which leads to silencing of tumor suppressor genes [40, 41]. In parallel, TFs also play instrumental roles in tumor development and metastasis [42–44]. Given the strong epigenetic basis of CRC, we next applied MethReg to the TCGA COAD-READ datasets, including 367 samples with matched DNA methylation, gene expression, and copy number alterations (CNAs). To account for potential confounding effects, we adjusted target gene expression values by CNA and tumor purity estimates [45] first, extracted the residuals, and then fitted the rlm.binary model to the residuals (see the “Methods” section for details).

We performed an unsupervised MethReg analysis, without selecting any CpGs *a priori*. First, we divided the CpGs into those in the promoter regions (within ± 2 kb regions around the transcription start sites (TSS)) or the distal regions (> 2kb from TSS). Next, we linked CpG sites in the promoter regions to genes with promoters that overlapped with them. On the other hand, CpG sites in the distal regions were linked to five genes upstream and five genes downstream within 1M bp. Alternatively, in the regulon-based approach, we also linked CpGs in either promoter or distal regions to genes regulated by TFs with binding sites close to the CpG (Figure 1). To more accurately model TF effects, we computed TF activity scores using the VIPER algorithm [33]. Stage-wise analysis using the rlm.binary model *residual target gene expression ∼ TF.activity + DNAm.group + DNAm.group × TF.activity* was then applied to the triplet datasets to identify triplets with significant *DNAm.group × TF.activity* effect, in which CpGs had significant effects on target gene expression by influencing TF activities.

Stage-wise analysis results showed the number of triplets with significant *DNAm × TF.activity* terms in the promoter, distal, and regulon-based analysis were 31, 52, and 47, respectively (Table 1, Supplementary Table S2 – S4). There was no overlap between the significant triplets obtained in these three analyses. Our results agreed well with the previous study in Wang et al. (2020) [30], which also observed only a small number of transcriptional regulations were mediated by DNA methylation. To additionally validate our findings, we compared the TFs in our significant triplets with the MeDReaders database [46], which manually curated information from human and mouse studies for 731 TFs which could bind to methylated DNA sequences. Our comparison showed that among the 18, 44, and 11 unique TFs identified in the promoter, distal, and regulon-based analyses, respectively, the majority of them were previously shown to interact with methylated DNA. More specifically, 12 (68%), 32 (73%), and 8 (73%) of the TFs from the promoter, distal and regulon-based analyses were previously shown to exhibit CpG methylation-dependent DNA-binding activities using functional protein arrays [14] or systematic evolution of ligands by exponential enrichment (SELEX) [16] (Supplementary Tables S2-S4).

**Table 1.** Stage-wise analysis results of TCGA COAD-READ datasets. The robust linear model target gene expression ∼ TF activity + DNAm + DNAm x TF was used to analyze CpG-TF-target gene triplets. Shown are significant triplets at 5% overall FDR level, and the unique CpGs, TFs, and targets in these triplets.

| analysis | unique triplets | unique CpGs | unique TFs | unique targets |
| --- | --- | --- | --- | --- |
| *Promoter analysis* |  |  |  |  |
| screening | 165 | 49 | 56 | 42 |
| confirmation | 31 | 29 | 18 | 25 |
| *Distal analysis* |  |  |  |  |
| screening | 2358 | 315 | 155 | 570 |
| confirmation | 52 | 43 | 44 | 44 |
| *Regulon-based analysis* |  |  |  |  |
| screening | 78 | 23 | 19 | 19 |
| confirmation | 47 | 18 | 11 | 6 |

Moreover, the TFs and target genes in the significant triplets identified by MethReg have also been previously associated with CRC. Figure 2 shows the most significant triplet among all the triplets considered in promoter and distal analysis. In this example, the transcription factor NFATC2 represses the target gene *HENMT1* in samples with high DNA methylation at cg00328227 but is relatively independent of target gene expression in samples with high DNA methylation. Therefore, DNA methylation enhances TF activities in this case. NFATC2 belongs to the NFAT family of transcription factors, which are known to regulate T cell activation and differentiation as immune cells invades the malignant tissues [47]. In particular, NFATC2 is a critical regulator in intestinal inflammation that promotes development of colorectal cancer, and expression levels of NFATC2 were found to be elevated in colorectal cancer patients [48, 49]. The MethReg prediction that NFATC2 represses *HENMT1* is consistent with the recent study that demonstrated higher expression of *HENMT1* is a favorable prognostic biomarker for colorectal cancer [50], and that NFATC2 is associated with tumor initiation and progression. The *HENMT1* gene encodes a methyltransferase that adds a 2’-O-methyl group at the 3’-end of piRNAs and was previously shown to be dysregulated in many cancers [51]. Our literature review showed the role of *HENMT1* is not well understood in colorectal cancer. Therefore, this example also shows MethReg can nominate plausible TF functions and targets that can be further studied experimentally. In contrast, without stratifying on DNA methylation levels, the direct TF-target association was only modest (Spearman correlation = - 0.033, P-value = 0.527).

Supplementary Figure 4 shows an example in which DNA methylation attenuates TF activity. In samples with low methylation level at cg02816729, the transcription factor TEAD3 down regulates target gene *SMOC2*. On the other hand, TEAD3 activity is relatively independent of target gene expression in samples with high methylation level at the CpG. TEAD3 belongs to the family of TEAD transcription factors, which also play critical roles in tumor initiation and progression in multiple types of malignancies including gastric, colorectal, breast, and prostate cancers [52]. Higher expression of TEADs, as well as association with poor patient survival, have been observed in many cancers including CRC [52, 53]. On the other hand, the target gene *SMOC2* was recently shown to be a favorable prognostic biomarker for better clinical outcomes in a large cohort of CRC patients [54]. Therefore, the MethReg prediction that transcription factor TEAD3 suppresses target gene *SMOC2* is consistent with the oncogenic role of TEAD3 and the favorable prognostic potential of *SMOC2*. Again, without stratifying on methylation levels, the TF-target association in all samples is only modest (Spearman correlation = 0.07, P-value = 0.183).

Interestingly, some of the TFs exhibited different modes of regulation depending on DNA methylation levels. In Supplementary Figure 5, the expression levels of target gene *BAHCC1* is increased by transcription factor POU3F4 in low methylation samples (P-value = 6.42×10^-4^) but is decreased by POU3F4 in high methylation samples (P-value = 1.31×10^-4^). Therefore, DNA methylation effectively increases the diversity of TF functions in this case. Without stratifying on DNA methylation, the TF-target association is not significant (Spearman correlation = 0.083, P-value = 0.112). POU3F4 belongs to the family of POU domain transcription factors, which are involved in different developmental processes and also recently found to be associated with the malignancy processes in different human tissues [55–59]. In addition, expression levels of the target gene *BAHCC1* were shown to be associated with survival times in different types of cancers such as melanoma [60], liver cancer [61] and pancreatic cancer [62].

Among other transcription factors in the top 10 triplets (Table 2), SNAI1 is a zinc-finger protein involved with inducing the epithelial-mesenchymal transition (EMT) process during which tumor cells become invasive with increased apoptotic resistance [63–65]. The MEIS1 homeodomain protein is a tumor suppressor and was observed to be down-regulated in colorectal adenomas [66]. Similarly, ISL2 is also a tumor suppressor and ISL2 loss increased cell proliferation and enhanced tumor growth in pancreatic ductal adenocarcinoma cells [67]. ETV4 belongs to a subfamily of the ETS family with well-known oncogenic properties [68]. In particular, ETV4 is over-expressed in colorectal tumor samples and higher ETV4 expression is associated with shorter patient survival time [69–75]. ZNF384 is a zinc finger protein that is found to be over-expressed in several cancers, including leukemia [76, 77], liver cancer [78], and colorectal cancer [79]. Finally, FOXL1 is an important regulator of the Wnt/APC/β-catenin pathway, which frequently activates event in gastrointestinal carcinogenesis [80–82]. Similarly, the target genes in the top 10 triplets have also been implicated previously in CRC and other cancers (Table 2). Moreover, among the significant triplets identified by MethReg, the majority of the CpGs, TFs, and target genes (78%, 79% and 75%, respectively) were also differentially methylated or differentially expressed between tumor and normal samples (Supplementary Tables 2-4). Taken together, these results demonstrated that MethReg is able to identify biologically meaningful signals from real multi-omics datasets.

**Table 2.** Top 10 most significant triplets identified by unsupervised MethReg analysis of TCGA COAD-READ samples. CRC = colorectal cancer.

| Triplet data |  |  |  | Annotations |  |  | DNAm x TF effect |  |  | TF-target |  | P-values for tumor vs. normal comparison |  |  | CRC literature |  |
| --- | --- | --- | --- | --- | --- | --- | --- | --- | --- | --- | --- | --- | --- | --- | --- | --- |
| ProbeID | position | TF | target gene | CpG-target distance | TF.role | DNAm.effect | Estimate | P-value | Adj. P-value | Correlation | P-value | CpG | TF | target gene | TF | target gene |
| *<u>promoter analysis</u>* |  |  |  |  |  |  |  |  |  |  |  |  |  |  |  |  |
| cg00328227 | chr1:<br>108661703 | NFATC2 | HENMT1 | -176 | Repressor | Enhancing | -0.663 | 0 | 0 | -0.033 | 5.27E-01 | 3.19E-08 | 2.53E-01 | 8.23E-08 | [47 ] [153] [49] | [51] [50] |
| cg25083481 | chr11:<br>1034989 | SNAI1 | MUC6 | 1727 | Activator | Attenuating | -0.413 | 1.77E-04 | 0 | 0.117 | 2.52E-02 | 8.98E-03 | 4.99E-10 | 1.97E-02 | [154] [63, 65] | [155] |
| cg02816729 | chr6:<br>168442419 | TEAD3 | SMOC2 | 1267 | Repressor | Attenuating | 0.323 | 7.77E-05 | 0 | 0.07 | 1.83E-01 | 2.96E-02 | 3.47E-02 | 7.32E-01 | [52] | [54] [156] |
| cg12751565 | chr1:<br>206946642 | NFATC2 | PIGR | -175 | Activator | Enhancing | 0.282 | 1.09E-09 | 0 | 0.315 | 8.67E-10 | 6.49E-03 | 2.53E-01 | 1.88E-11 | [47 ] [153] [49] | [157] |
| cg05503887 | chr11:<br>34620378 | MEIS1 | EHF | -713 | Activator | Attenuating | -0.268 | 7.24E-05 | 0 | 0.347 | 1.10E-11 | 6.17E-01 | 1.84E-08 | 1.16E-02 | [66] | [158] |
| cg13542964 | chr4:<br>10457043 | ETV4 | ZNF518B | 381 | Activator | Enhancing | 0.217 | 2.08E-04 | 0 | 0.023 | 6.59E-01 | 1.31E-06 | 1.32E-14 | 1.77E-02 | [68, 72] | [159] |
| *<u>distal analysis</u>* |  |  |  |  |  |  |  |  |  |  |  |  |  |  |  |  |
| cg14043104 | chr13:<br>74075149 | ISL2 | KLF12 | -80092 | Activator | Attenuating | -0.222 | 1.03E-04 | 0 | 0.232 | 7.60E-06 | 3.89E-02 | 3.96E-02 | 1.20E-03 | [67] | [160] |
| cg09217215 | chr11:<br>31808034 | ZNF384 | PAX6 | 9925 | Repressor | Enhancing | -0.398 | 6.90E-07 | 1.48E-04 | -0.028 | 5.94E-01 | 7.64E-03 | 2.15E-01 | 1.68E-02 | [76] [77] | [161] |
| cg11871337 | chrX:<br>56991382 | FOXL1 | SPIN3 | 4443 | Activator | Attenuating | -0.294 | 2.18E-07 | 2.68E-04 | 0.183 | 4.32E-04 | 7.26E-01 | 9.95E-01 | 8.21E-07 | [80] [81] | [162] |
| cg04665204 | chr17:<br>81453398 | POU3F4 | BAHCC1 | 57922 | Dual | Invert | -0.37 | 6.80E-07 | 2.91E-04 | 0.083 | 1.12E-01 | 2.69E-05 | 1.47E-06 | 2.38E-02 | [55] [59] | [62] [61] |

#### Comparative analysis of MethReg (in unsupervised mode) with alternative approaches

Encouraged by the consistency of MethReg results with recent colorectal cancer literature, we next performed a comparative analysis of MethReg with several alternative approaches. As evidence from large-scale analyses that a substantial number of TFs can interact with methylated DNA have only become available in recent years, few studies have analyzed DNA methylation, TF binding sites, and gene expression data simultaneously using large multi-omics datasets until the past year.

We considered a total of three alternative approaches: the recent studies by Wang et al. (2020) [30] and Liu et al. (2019) [83], as well as the conventional approach of directly correlating DNA methylation with gene expression. In Wang et al. (2020), a rewiring score constructed based on TF-target gene correlations in high and low methylation samples was proposed to identify methylation-sensitive TFs in different cancers using TCGA data. Similarly, using a conditional mutual information-based approach, Liu et al. (2019) identified CpG-TF-target triplets in which the TF-target regulation circuit is dependent on CpG methylation levels in different cancers using TCGA data. In both of these studies, sample permutations were used to estimate P-values for test statistics based on rewiring scores or conditional mutual information. Conceptually, compared to these previous studies, MethReg makes several distinct contributions: (1) MethReg analysis is comprehensive. While both previous works [30, 83] analyzed only promoter CpG methylation, MethReg analyzes methylation CpGs at both promoter and distal (enhancer) regions. The identification of distal regulatory elements is crucial as they are often not well defined. (2) MethReg analysis is flexible. In addition to the example databases we have described here, MethReg is capable of incorporating any user-specified ChIP-seq or regulons database (Supplementary Table 1), including those that are tissue- or disease-specific. The power and potential of MethReg grows as more knowledge on TF regulation becomes available. (3) MethReg analysis is rigorous. Robust linear models are implemented to carefully model outlier samples in RNA-seq data. Further, with the permutation approaches used in the previous studies, it can be challenging to adjust for covariate information that is important for analyzing human datasets, for which confounding variables such as tumor purity are often significant contributors to both DNA methylation and gene expression. In contrast, MethReg’s regression-based approach makes it easy to adjust for potential confounding variables. (4) Most importantly, while previous studies provided analysis results for TCGA cancer datasets, until now no software has been made available to the research community. This study provides an open-source software, MethReg, which implements careful bioinformatics and statistical analysis, to make it possible for researchers to analyze datasets beyond TCGA. In addition to providing estimated individual and joint regulatory effects of CpG methylation and TFs, MethReg also annotates the potential regulatory roles of TFs and CpG methylation, as well as providing a rich suite of figures for visualizing analytical results.

Next, we performed a systematic comparison of MethReg analysis results and results from other approaches empirically using the TCGA COAD dataset. Because both previous studies analyzed TF gene expressions, we performed additional MethReg promoter analysis that model the TF effect based on TF gene expression, instead of TF activities. More specifically, to compare with results from Wang et al. (2020), which identified 3,244 TF-target gene pairs, we applied the robust linear model described above to the corresponding triplets (average promoter DNA methylation, TF gene expression, and target gene expression). Our results showed that for a majority (81.23%, n = 2613) of the significant TF-target pairs from Wang et al. (2020), the corresponding triplets also had significant DNAm × TF effect in the rlm.binary model from MethReg. Moreover, classification for promoter methylation effects on TFs in Wang et al. (2020) and MethReg agreed very well (kappa statistic = 0.975, P-value = 0) (Supplementary Table 5).

Similarly, we also fitted our rlm.binary model described above to the 47,029 triplets identified in Liu et al. (2020) for the TCGA COAD dataset. However, we observed less agreement between our significant results and those from Liu et al. (2019), as only 5,321 (11.3%) of the 47,029 triplets had significant DNAm x TF p-values by MethReg. The overlap between Liu et al. (2019) with Wang et al. (2020) was also very low, with only 8 TF-target associations identified by both studies. This discrepancy might be due to the differences in methodologies. The conditional mutual information approach used by Liu et al. (2019) detects any general associations which can be non-monotonic, while the robust linear model MethReg used and the rewiring score Wang et al. (2020) used mainly detects linear and monotonic associations in TF and target genes that are dependent on CpG methylation.

Finally, we compared MethReg promoter analysis results with the conventional approach of correlating methylation-target gene directly. More specifically, Spearman correlations were computed for each promoter CpG and target gene in the COAD-READ dataset. The correlation between rankings of P-values based on methylation-target gene correlation and *DNAm × TF* interaction effects in the MethReg model is a significant but modest association (Spearman correlation = 0.0533, P-value = 2.2 × 10^-16^), indicating these two approaches are identifying many different CpGs. This is not surprising, however, because the MethReg model identifies CpG methylation that influences the target gene by regulating TF activities (Figure 5B), instead of influencing the target gene directly (Figure 5A).

**Figure 5.**
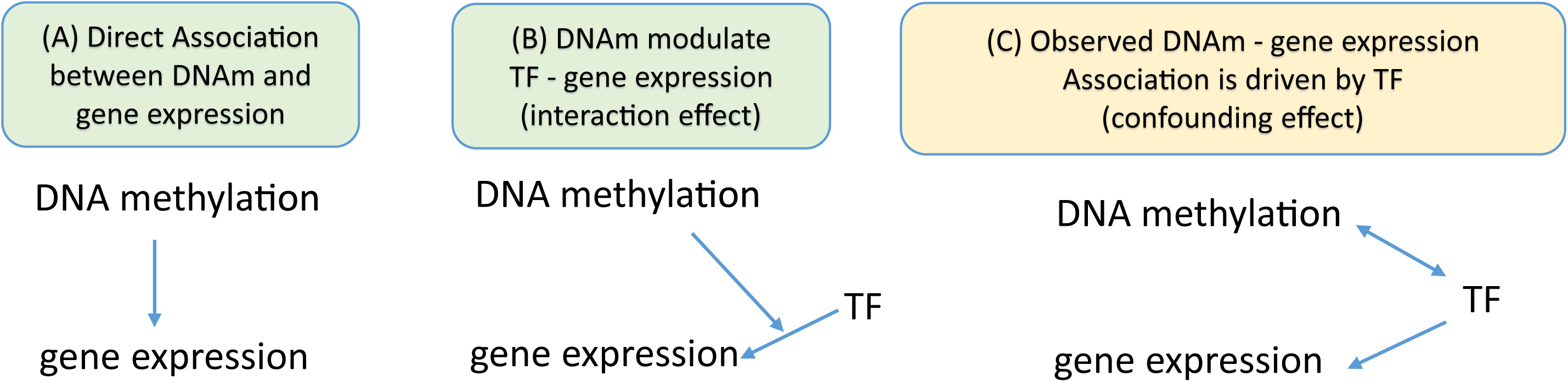
Direct, indirect, and confounding effects in gene regulation.

While MethReg is not designed to identify direct methylation-target associations, it can be useful for identifying those methylation-target correlations likely driven by TF effects (Figure 5C). In this case, TF activities are associated with DNA methylation levels, and TF also regulates gene expression independently of DNA methylation (Fig 5C). In statistics, TF represents a confounding effect in the methylation-target gene association. For example, Supplementary Figure 6 shows an example from the analysis of triplets in Wang et al. (2020) described above, where the observed correlation between promoter DNA methylation and target gene expression is highly significant (Spearman correlation = 0.238, P-value = 4.79 × 10^-5^, FDR = 2.81 × 10^-4^). However, the result of fitting the MethReg model indicated that for this triplet, neither *DNAm* nor *DNAm* × *TF* terms were significant (P-values = 0.395 and 0.477, respectively), but *TF* was highly significant (P-value = 5.75 × 10^-5^). Therefore, the target gene expression is likely driven mainly by the TF EBF1, a tumor suppressor with prognostic value for CRC [84], and not by DNA methylation, even though we observed a highly significant methylation-target gene expression association.

### Case study: analysis of ROSMAP Alzheimer’s disease dataset

#### A supervised MethReg analysis

In this section, we demonstrate supervised MethReg analysis using an Alzheimer’s disease (AD) dataset collected by the ROSMAP study [85]. In contrast to unsupervised analysis which tests triplets involving all CpGs, in a supervised analysis, we only test triplets involving differentially methylated CpG sites (DMS), typically obtained from EWAS studies. To study AD associated DNA methylation changes in the brain, we recently performed a meta-analysis of over 1,000 prefrontal cortex brain samples from four large brain studies [6, 86–88], and identified 3,751 significant CpGs at 5% FDR [89]. To help understand the regulatory roles of these DMS, we applied MethReg to analyze matched DNA methylation and gene expression profiles measured on prefrontal cortex brain samples from 529 independent subjects in the ROSMAP dataset.

To illustrate the versatility of the MethReg analysis pipeline, we used alternative databases to analyze the ROSMAP dataset, compared to the analysis pipeline for the TCGA COAD-READ samples. More specifically, to map TF binding sites, instead of the JASPAR2020 database [30] here we used the ReMap database [26], which contains a large collection of regulatory regions obtained using genome-wide DNA-binding experiments such as ChIP-seq. In particular, the ReMap human atlas included binding regions for 1,135 transcriptional regulators. Also, to analyze these brain samples, instead of the pan-cancer regulons, we used the brain-specific TF regulons included in the ChEA3 [90] software website, along with TF activity scores estimated by Gene Set Variation Analysis (GSVA) [34].

More specifically, we first adjusted methylation and gene expression values separately by potential confounding effects, including age at death, sex, batch effects, and markers of different cell types. Next, we computed TF activities using GSVA method [34], which is an alternative method to VIPER [33] for computing enrichment scores of each TF, by comparing enrichment in target gene expressions for a TF (its regulons) with expression levels of background genes.

At 5% false discovery rate, MethReg identified 1, 20, and 103 triplets which included 1, 16, and 53 unique TFs that interact with DMS to influence target gene expressions in promoter, distal, and regulon-based analyses, respectively. A comparison with the MeDReaders database [46] shows more than half (58.6%, 41 out of 70) of these TFs were previously shown to interact with methylated DNA sequences (Supplementary Tables 6 and 7). In Table 3, many of the TFs and target genes in the top 10 triplets were previously implicated in AD pathology. For example, in the most significant triplet (Table 3, Supplementary Figure 7), the transcription factor SPI1 (PU.1) is a master regulator in the AD gene network [91, 92]. SPI1 is critical for regulating the viability and function of microglia [93], which are resident immune cells of the brain. Microglia functions as primary mediators of neuroinflammation and phagocytoses amyloid-beta peptides accumulated in AD brains [94]. Using transgenic mouse models for AD, and comparing with gene-level variations in recent human AD GWAS meta-analysis [95], Salih et al. (2019) [96] recently showed the target gene *LAPTM5* (lysosome-associated protein, transmembrane 5) belongs to an amyloid-responsive microglial gene network, and predicted *LAPTM5* to be one of four new risk genes for AD. Moreover, *LAPTM5* was also shown to be a member of human microglia network in AD in multiple gene expression studies [97, 98]. Intriguingly, SPI1 and *LAPTM5* belonged to the same transcription co-expression network [96], and in mouse microglial-like BV-2 cells, results from ChIP-seq experiment showed SPI1 binds to the regulatory region of *LAPTM5* [99], consistent with the MethReg prediction that *LAPTM5* is regulated by SPI1. Interestingly, SPI1 appears to have dual roles in this case, depending on DNA methylation levels at cg17418085, which is located in the gene body of *LAPTM5*. More specifically, SPI1 upregulates the target gene *LAPTM5* in samples with low methylation at cg17418085 but down-regulates the target gene when DNA methylation is high, suggesting DNA methylation and the TF might have compensatory mechanisms that control gene expression at this locus (Supplementary Figure 7).

**Table 3.** Top 10 most significant triplets identified by supervised MethReg analysis of ROSMAP Alzheimer’s disease (AD) dataset.

| Triplet data |  |  |  | Annotation |  |  | DNAm x TF effect |  |  | TF-target |  | AD literature |  |
| --- | --- | --- | --- | --- | --- | --- | --- | --- | --- | --- | --- | --- | --- |
| ProbeID | position | TF | target gene | CpG-target distance | TF.role | DNAm.effect | Estimate | P-value | Adj. P-value | Correlation | P-value | TF | target gene |
| *<u>promoter analysis</u>* |  |  |  |  |  |  |  |  |  |  |  |  |  |
| cg17418085 | chr1:<br>31229122 | SPI1 | LAPTM5 | 1,543 | Dual | Invert | -7.177 | 7.36E-12 | 3.06E-07 | -0.011 | 8.05E-01 | [92, 93] | [96] |
| *<u>distance-based analysis</u>* |  |  |  |  |  |  |  |  |  |  |  |  |  |
| cg11556846 | chr2:<br>200468728 | ESR1 | SATB2 | -132,738 | Dual | Invert | -7.266 | 3.03E-07 | 7.80E-03 | -0.058 | 1.82E-01 | [111, 113] | [163] [164] |
| cg00153919 | chr16:<br>88859944 | CEBPD | PIEZO1 | -8,324 | Repressor | Enhancing | -8.122 | 4.92E-07 | 9.14E-03 | -0.044 | 3.10E-01 | [117, 118] | [165] |
| cg08824847 | chr11:<br>35052388 | NR3C1 | PDHX | 115,011 | Activator | Enhancing | 4.383 | 6.13E-07 | 1.02E-02 | 0.06 | 1.66E-01 | [101] [103] | [104, 105] |
| cg08760493 | chr4:<br>109994039 | TCF12 | SEC24B | -360,887 | Dual | Invert | 8.120 | 1.26E-06 | 1.49E-02 | 0.033 | 4.44E-01 | [119, 120] | [166] |
| cg05715492 | chr7:<br>98991138 | SRF | ARPC1B | 19,265 | Repressor | Attenuating | 11.581 | 1.34E-06 | 1.49E-02 | 0.017 | 7.00E-01 | [122] | [167] |
| cg09316954 | chr16:<br>67687754 | TCF12 | CARMIL2 | 8,931 | Repressor | Attenuating | 8.376 | 2.28E-06 | 2.24E-02 | -0.07 | 1.08E-01 | [119, 120] | [168] |
| cg21155834 | chr2:<br>149282209 | GABPA | ORC4 | -503,061 | Activator | Enhancing | 6.407 | 2.75E-06 | 2.41E-02 | 0.045 | 2.98E-01 | [124] | [169] |
| cg13819552 | chr9:<br>95799870 | TCF12 | ZNF484 | -159,565 | Activator | Enhancing | 7.989 | 2.92E-06 | 2.41E-02 | 0.039 | 3.71E-01 | [119, 120] | [170] |
| cg21535772 | chr2:<br>171679906 | NFE2L2 | AC007405.4 | 52,282 | Dual | Invert | 7.065 | 3.10E-06 | 2.41E-02 | -0.026 | 5.47E-01 | [125] [171] | [172] |

The triplet cg08824847- NR3C1- PDHX is an example in which methylation at cg08824847 appears to enhance activation of the target gene *PDHX* by NR3C1 (Table 3, Supplementary Figure 8). NR3C1 is the glucocorticoid receptor (GR) which can act as a transcription factor that binds glucocorticoid responsive genes to activate their transcriptions, or as a regulator for other TFs. NR3C1 regulates downstream processes such as glycolysis [100] and was observed to be dysregulated in AD [101–103]. The target gene *PDHX* encodes the component X of the pyruvate dehydrogenase complex (PDH), which is involved in the regulation of mitochondrial activity and glucose metabolism in the brain that is critical for neuron survival [104, 105]. Low levels of cerebral glucose metabolism often proceed with the onset of Alzheimer’s Disease and have been proposed as a biomarker of AD risk [106–109]. Previously it was also shown that along with other regulators, glucocorticoids can regulate the efficacy of PDH [100, 110]. The MethReg prediction that methylation at cg08824847 enhances activation of *PDHX* by NR3C1 is consistent with these previous findings that demonstrated lower levels of *PDHX* in AD samples, and with results from our previous large meta-analysis of DNA methylation changes in AD, which discovered cg08824847 to be hypo-methylated in AD samples across all four analyzed brain samples cohorts and has a significant negative association with AD Braak stage even after multiple comparison correction [89].

Among the other TFs in the top 10 triplets (Table 3), ESR1 is estrogen receptor alpha, one of two subtypes of estrogen receptor. Genetic polymorphisms of ESR1 have been associated with risk of developing cognitive impairment in older women [111–115], as well as faster cognitive decline in women AD patients [116]. Induced by chronic inflammation in AD, CEBPD is associated with microglia activation and migration [117, 118]. TCF12 is a member of the basic helix-loop-helix (bHLH) E-protein family and plays important roles in developmental processes such as neurogenesis, mesoderm formation, and cranial vault development. Recently, TCF12 was predicted to be affected by SNP rs10498633 [119], a top AD-associated SNP identified in the IGAP AD meta-analysis study [120]. Moreover, TCF12 also belongs to a herpesvirus perturbed transcription factor regulatory network that is implicated in AD [121]. SRF is serum response factor, responsible for regulating the smooth muscle cells and blood flows in the brain, which are important for a blood vessel’s ability to remove amyloid beta peptides accumulated in AD. Compared with healthy individuals, SRF was found to be four times higher in AD patients [122, 123]. GABPa belongs to the ETS family of DNA-binding factors and is a master regulator of multiple important processes including cell cycle control, apoptosis, and differentiations. Using evolutionary analysis and ChIP-seq experiments, Perdomo-Sabogal et al. (2016) [124] linked GABPa to several brain disorders including AD, autism, and Parkinson’s disease. Finally, NFE2L2/NRF2 is another master regulator and regulates genes involved in response to oxidative stress and inflammation. Motivated by the encouraging therapeutic effect of NRF2 on AD pathology in animal models and cultured human cells [125–127], modulation of NRF2 pathway has recently been proposed as a strategy for AD drug development [125]. Similarly, Table 3 shows the target genes in these top 10 triplets identified by MethReg are also highly relevant to Alzheimer’s disease. Taken together, these results demonstrated MethReg is also capable of identifying biologically meaningful regulatory effects of DNA methylation in other complex diseases, such as the AD brain DNA methylation data we used here, where association signals are expected to be much weaker than those observed in cancers.

### Comparative analysis of MethReg (in supervised mode) with alternative approaches

To compare the performance of MethReg in supervised mode with currently available alternative tools, we next analyzed the ROSMAP dataset using the ReMapEnrich R package [26], which identifies regulators with binding sites enriched in user supplied regions. Several other tools such as LOLA [21] and ChIP-Enrich [128] perform similar analyses as ReMapEnrichR, but here we chose ReMapEnrichR because it uses the same ReMap database as MethReg for the ROSMAP dataset analysis. More specifically, for the ReMapEnrichR analysis, we also used the locations of the 3,751 AD-associated CpGs (these are the DMS) from previous AD meta-analysis as the input. The results showed that ReMapEnrich identified 143 TFs with binding sites enriched in the DMS, among which a substantial number (n = 28, 20%) were also identified by MethReg (Supplementary Table 8). These 28 TFs included many well-known regulators for AD such as TCF12, SPI1, NR3C1, CEBPB, GABPA and others. On the other hand, 115 and 32 TFs were uniquely identified by ReMapEnrich or MethReg, respectively.

Although many of these significant TFs have been previously implicated in AD pathology, their specific roles in transcription regulation and the identification of their targets in AD remain to be investigated. Notably, currently available tools such as ReMapEnrich only identify the TFs but do not consider CpGs or provide detailed information on the relevant target genes. In contrast, MethReg fills this critical gap by nominating plausible TF-target associations that are mediated by DNA methylation. Therefore, MethReg analysis which leverages additional gene expression data and provides more comprehensive information on transcription regulation for the TFs complements existing approaches.

## Discussion

To evaluate the role of DNA methylation in gene regulation, we developed the MethReg R package. MethReg provides a systematic approach to dissect the variations in gene transcription into three different modes of regulation, which are direct effects by methylation and TF individually, and the synergistic effect from both DNA methylation and TF. By additionally modeling DNA methylation variations, MethReg complements existing approaches that analyze TF and target gene expression alone. In doing so, MethReg uncovers TF-target gene relations that are present only in samples with high (or low) methylation levels at the CpG that mediates TF activities. On the other hand, compared to approaches that analyze DNA methylation and TFBS data alone, MethReg analysis also reduces the noisiness in TF binding site predictions by additionally modeling target gene expression data. Compared to the conventional approach of directly correlating DNA methylation with gene expression, MethReg can be useful for identifying methylation-gene expression associations which are likely driven by TFs instead of methylation due to confounding effects by TFs. Computationally, MethReg is efficient. The unsupervised analysis of TCGA COAD-READ datasets which considered all CpGs on the Illumina array took 5, 37, and 14 minutes for the promoter, distal, and regulon-based analyses, respectively, using a single Linux machine with 64 GB of RAM memory and Intel Xeon W-2175 (2.50 GHz) CPUs with 4 cores for parallel computing (Supplementary Table 9).

Because of current limitations in technology, directly measuring DNA methylation, TF binding and target gene expression in high throughput is still a difficult task, especially for a large cohort of primary tissue samples. Therefore, computational approaches are needed to prioritize regulatory elements in gene transcription. To this end, a main computational challenge is the accurate assessment of TF activities. Many integrative studies have used TF gene expression data, which are often widely available, as surrogate measurements for TF activities. However, the abundance of TF expression does not necessarily correspond to more TF binding events, which needs to be confirmed by cell type specific ChIP-Seq experiments. On the other hand, TF binding events are sometimes non-functional and might not lead to changes in gene expression [24, 25]. To this end, we implemented the option to model TF effects based on VIPER [33] or GSVA [34] estimated TF protein activities in MethReg. Both of these methods assume that collectively, the target genes of a TF represent an optimal reporter of its activity and estimate TF activity based on enrichment of its target genes expressions compared to background genes. Both VIPER and GSVA approaches have been widely used for modeling protein activities using gene expression datasets.

While the motivation of MethReg is to rank significant and functional DNA methylation changes identified in EWAS, a useful by-product of this analysis is the identification of enhancers, which are often located several hundreds of kilobases (kb) away from the target gene, where TFs bind and interact with DNA methylation to activate gene expression by looping DNA segments [129]. Growing evidence indicates that in addition to promoter methylation, DNA methylation at enhancers also plays an equally or more important role in activating gene expression [130]. Active cancer-specific enhancers are typically hypomethylated at CpG sites [10, 131, 132] in open chromatin regions free of nucleosomes [133, 134]. In many cases, hypermethylation of CpG sites can interfere with TF binding and lead to decreased enhancer activities in various cancers [9, 135]. It has been observed that TF activities often correlate with levels of demethylation at enhancer regions and subsequent target gene expressions [135–137].

Although recently many cis-regulatory regions have been identified using genomic and epigenomic data [136, 138], assigning these candidate enhancers to target genes on a genome-wide scale remains challenging and is currently an active area of research. A recent study [139] compared several published computational approaches for enhancer-gene linking using a collection of experimentally derived genomic interactions. It was shown that the best-performing method, TargetFinder [140], is only modestly better than the baseline distance-based approach, and the authors suggested further improvement in current computational methods in this area is needed. Although many of these computational methods leverage various information from histone marks, chromatin accessibility and interaction, TF binding models and gene expression levels, few if any also model DNA methylation at candidate enhancer regions. As many recent studies suggested substantial crosstalk between DNA methylation and other regulatory elements such as transcription factors [14, 16], we developed MethReg to specifically estimate and evaluate the regulatory potential of DNA methylation for interacting with candidate TFs at both promoter and distal regions. In particular, MethReg links target genes with CpGs in distal regions using two alternative approaches: linking to a fixed number of nearby genes or by using annotations in regulon databases (Figure 1). Indeed, among significant triplets identified in MethReg analysis of ROSMAP dataset, a substantial number of the CpGs (6 and 22 in distal and regulon-based analyses, respectively) were located in brain-specific enhancer regions annotated in the EnhancerAtlas 2.0 database [141] (Supplementary Tables 6-7). Notably, the power and potential of MethReg will also grow as more knowledge on TF regulatory activities are accumulated and new ChIP-seq and TF regulon databases become available.

The aim of MethReg is to prioritize functional elements and to generate useful testable hypothesis for subsequent mechanistic studies. For the DNA methylation changes identified by MethReg which couple with TF activities, additional experimental studies are needed to determine whether the changes in methylation are causing or are caused by TF activities. Nevertheless, even if the DNA methylation changes are passive markers that accumulated as result of TF binding (or lack of binding), they can still be useful in the clinics as biomarkers. Currently, many of the large DNA methylation datasets are measured using methylation arrays because of their lower cost and the simplicity in benchwork and analysis. MethReg has been tested successfully on microarrays and can also be easily extended to analyze large cohorts of samples measured using high throughput sequencing such as WGBS or RRBS. In addition to transcription factors, MethReg can also be applied to analyze other types of chromatin proteins including histones which are known to crosstalk with DNA methylation [142–144]. Finally, MethReg can further be extended to incorporate TF-target associations based on spatial enhancer-gene linking when 3D chromatin and genetic interaction data on primary cells become available.

## Conclusions

We have presented an integrative analysis and annotation software, MethReg, which has several critical roles. First, given the large number of DMS identified from EWAS, supervised MethReg analysis can be used to analyze CpG-TF-target gene triplets involving the DMS, to identify and prioritize important CpG methylations that influence target gene expressions by interacting with TFs that bind in proximity. Second, MethReg annotates CpG methylation and the TFs that bind in close proximity with their regulatory roles (i.e., activator or repressor TFs, DNA methylation that attenuate or enhance TF effects). Third, because some TFs affect target gene expression only in samples with high methylation levels (or only in samples with low methylation levels), MethReg can help uncover TF-target gene associations that are not obvious in an analysis that uses all samples. Finally, for a particular target gene, MethReg partitions the variances in gene regulation into direct impact by methylation, direct impact by TF, or joint impact of both methylation and TF, which allows MethReg to identify methylation-target gene associations that are likely driven by TF instead of DNA methylation. Using two case studies in colorectal cancer and Alzheimer’s disease, we have shown the power of MethReg to uncover biologically relevant transcriptional regulation in both diseases which have vastly different biology. Open source software scripts, along with extensive documentation and example data for MethReg are freely available from the Bioconductor repository. We hope MethReg will empower researchers to gain a better understanding of the important regulatory roles of CpG methylation in many complex diseases.

## Methods

### Unsupervised MethReg analysis of TCGA COAD-READ samples

#### Datasets

Level 3 TCGA COAD and READ datasets, including genomic profiles in copy number alterations (CNAs; Gene Level Copy Number Scores), gene expressions (HTSeq - FPKM-UQ values from RNA-seq), and DNA methylation levels (beta values from 450k Illumina arrays) were downloaded from the NCI’s Genomic Data Commons (GDC) using TCGAbiolinks R package (version 2.17.4) with functions GDCquery, GDCdownload, and GDCprepare [145]. These level 3 datasets were previously pre-processed and normalized as described in https://docs.gdc.cancer.gov/Data/Bioinformatics_Pipelines. We included only primary tumors samples in our analysis. Tumor purity estimates were downloaded from Supplementary Data 1 of Aran D et al. (2015) [45]. We included 367 samples with DNA methylation, gene expression, and copy number alterations (CNAs) profiles. To compute TF activity scores, we used the pan-cancer regulon database from Garcia-Alonso et al. (2019) [29], downloaded from https://github.com/saezlab/dorothea/tree/deprecated/data/TFregulons/consensus/Robjects_VIPERformat/pancancer

#### Pre-processing data

First, we removed genes and CpGs with little expression variations across samples. More specifically, from a total of 19,456 genes with matched DNA methylation, gene expression, and CNA values, we removed 2,315 genes with zero counts in more than 50% of the samples in RNA-seq data. For each CpG, we computed inter-quartile range (IQR) of the beta values and removed CpGs with IQR less than 0.2. In addition, to avoid confounding by TF effect (Figure 5C), we focused the analysis on those CpG-TF-target gene triplets where CpG was associated with the target gene (Wilcoxon test p-value less than 0.05), but was not associated with TF (Wilcoxon test p-value greater than 0.05).

Next, for each gene, the get_residuals function in MethReg was used to remove potential confounding effects in gene expression data by fitting a linear model which included log_2_ transformed gene expression as the outcome variable, and CNA and tumor purity estimates as predictor variables. The residuals obtained from this model were then used for subsequent analyses. To estimate TF activity scores, we used the VIPER algorithm [33], which is implemented in function run_viper in R package dorothea, along with the pan-cancer regulons described above.

#### Creating triplet datasets

In the promoter analysis, the function create_triplet_distance_based with parameter target.method = “genes.promoter.overlap” were used to link CpGs in the promoter regions (within ± 2 kb regions around the transcription start sites (TSS)) to the corresponding target genes. Similarly, in distal analysis, the function create_triplet_distance_based with parameters target.method = “nearby.genes” and target.num.flanking.genes = 5 were used to link CpGs in the distal region (beyond ± 2 kb regions around the TSS) to five genes upstream and downstream of the CpG. Alternatively, in regulon-based analysis, the function create_triplet_regulon_based was used to link CpGs to the TF-target gene pairs in the pan-cancer regulons. In all three analyses, we obtained TF binding sites information from the JASPAR2020 [27] database and linked CpGs to TFs with binding sites within 250 bp distance using *motifmatchr* R package (version 1.10.0). The CpG-TF pairs were then combined with CpG-target gene pairs (promoter and distal analyses) or TF-target pairs (regulon-based analysis) to obtain CpG-TF-target triplets. In all analysis, we specified the parameter max.distance.region.target = 10^6 to obtain triplets for which the CpG is within 1M bp distance from the target gene. These analyses were performed using hg38 genome assembly coordinates.

#### Statistical analysis

Given CpG-TF-target gene triplets from “Creating triplet datasets” section, and DNA methylation, estimated TF activities, and residual gene expression data from “Pre-processing data” section, the function interaction_model was then used to fit robust linear statistical model *residual.target.gene.expression ∼ TF + DNAm.group + DNAm.group × TF.* Within the interaction_model function, the robust linear model analysis is implemented by rlm function from the MASS R package, with parameter psi = psi.bisquare (bisquare weighting) [146], which gives outlier gene expression values reduced weight. After p-values for the interaction term *DNAm.group* x *TF* was computed for each triplet, we next performed stage-wise analysis by setting stage.wise.analysis = TRUE in interaction_model function. The stage-wise analysis in *MethReg* is implemented by calling stageRTx function of stageR R package [37]. For each CpG, we grouped all triplets associated with it and used the smallest p-value for *DNAm.group × TF* to represent the CpG. These smallest p-values for each CpG were then used to define values for pScreen in function stageRTx. The parameter pConfirmation was defined as *DNAm.group* x *TF* p-values for each triplet. In the screening step, the null hypothesis that any of the individual triplets mapped to a CpG had a significant *DNAm.group × TF* effect was tested. In the confirmation step, all the triplets associated with significant CpGs selected in the screening step were tested while controlling FWER [37].

#### Annotating roles of TFs and effects of CpG methylations

For triplets with significant *DNAm.group × TF*, we next annotated roles of TF and CpG methylations in gene regulation (Figure 3). To this end, the function stratified_model fits the robust linear model *residual.target.gene.expression* ∼ *TF* in samples with high or low methylation levels at the relevant CpG separately, to obtain p-values for TF effect (pval.tf.high and pval.tf.low). Let the most significant p-value among the two p-values be pval.most.sig. A TF is annotated only if pval.most.sig < 0.001 (that is pval.tf.high < 0.001 or pval.tf.low < 0.001). If the estimated TF effect (i.e., slope) corresponding to the most significant p-value (pval.most.sig) is positive, the TF is annotated as an activator. If the estimated TF effect corresponding to pval.most.sig is negative, the TF is annotated as a repressor. To annotate CpG methylation effect, if pval.most.sig is from the model fitted to high methylation samples, the CpG is labeled as enhancing; otherwise the CpG is labeled as attenuating. If pval.tf.high < 0.001 and pval.tf.low < 0.001, and the TF effects (i.e., slopes) in high and low CpG methylation samples have opposite signs, then the TF is annotated to have dual roles and the CpG is labeled to have invert effect.

#### Tumor vs. normal samples analysis

For each significant triplet from the stageR analysis described above, we additionally assessed differential methylation of the CpGs and differential expressions of the TFs and target genes in tumor vs. normal samples. To this end, 21 adjacent normal samples with matched gene expression (HTSeq - FPKM-UQ values from RNA-seq) and DNA methylation levels (beta values from 450k Illumina arrays) were downloaded from NCI’s Genomic Data Commons (GDC) using TCGAbiolinks R package (version 2.17.4) using functions GDCquery, GDCdownload *and* GDCprepare [145]. To assess the statistical significance of differential methylation levels at CpGs (or differential gene expression levels at TF and target genes) in tumor vs. normal samples, we used two-tailed Wilcoxon test.

### Simulation study

The *rlm.binary.en* method estimates P-values for each triplet using empirical null distribution [38, 147], which is a normal distribution with empirically estimated mean 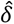 and standard deviation 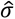. There are five steps:

(1) For each triplet, fit rlm.binary model to obtain *t-*statistic and degrees of freedom (*df*) corresponding to *DNAm.group × TF* effect.
(2) Convert the t-statistic to a z-score. For example, let *t_i_* be the *t-*statistic for triplet *i*, the corresponding z-score can be obtained by *z_i_* = Φ^−1^(*F_d_*(*t_i_*,*d_f_*)) where Φ and *F_d_* are distribution functions for standard normal distribution and *t* distribution with *df* degrees of freedom.
(3) Pool the z-scores for all triplets tested in an analysis and calculate their median value (*m*). Subtract *m* from the z-scores so that they have median of 0.
(4) Given the median-centered z-scores, use the locfdr package, estimate location (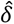) and scale (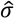) parameters of the empirical null distribution.
(5) Calculate standardized z-scores, 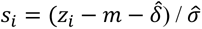, and compute the *P*-value for each triplet as *p_i_* = 1 − Φ(*s_i_*).

In triplets simulated under the scenario β = 0, target gene expressions did not depend on TF expression levels, so the genes simulated under this simulation scenario were the null triplets. Type I error rate for each model was computed as the number of triplets declared to be significant by the model when β = 0, divided by the total number of null triplets. P-values less than 0.05 for each method were considered to be significant. On the other hand, power was computed using triplets simulated under the scenarios where *β* = 1, .., 9. For each of the 7 compared methods (Supplementary Figure 1), power was estimated as the number of triplets with P-values less than 0.05 for each compared method, divided by the total number of simulated triplets (i.e. 1,000). The pROC R package was used to compute area under ROC curves.

### Comparative analysis with alternative approaches

For the comparison with Wang et al. (2020) [30], we downloaded data from the DMTDB database (http://bio-bigdata.hrbmu.edu.cn/DMTDB; accessed on June 5, 2020). The file “COAD_DMTD” contained 3,244 COAD DNA methylation dependent TF-target gene pairs, where target gene expressions were regulated by the TF and average DNA methylation at promoter region (± 2k bp) of the target gene. For the comparison with promoter analysis results from Liu et al. (2019) [83], we downloaded promoter CpG methylation-dependent TF-target associations from file “COAD.txt” in Supplementary Data S1 of Liu et al. (2019). We included 42,590 triplets with non-masked CpGs [148] in our analysis.

### Analysis of ROSMAP Alzheimer’s disease dataset

#### Datasets

The ROSMAP dataset, which included samples from the Religious Order Study (ROS) and the Memory and Aging Project (MAP) [85], included 529 samples with matched DNA methylation and gene expression data [6]. More specifically, normalized FPKM (Fragments Per Kilobase of transcript per Million mapped reads) gene expression values and Illumina HumanMethylation 450K beadchip files (.idat format) for the ROSMAP study were downloaded from the AMP-AD Knowledge Portal (Synapse ID: syn3388564). Brain-specific regulons (file “brain.TFs.gmt”) were downloaded from ChEA3 software website https://maayanlab.cloud/chea3/ (accessed Oct 10, 2020).

#### Pre-processing datasets

DNA methylation pre-processing included the removal of CpG probes with detection P-value > 0.01 across all the samples in the cohort, and CpG probes in which a single nucleotide polymorphism (SNP) with minor allele frequency (MAF) ≥ 0.01 was present in the last 5 base pairs of the probe. The quality controlled methylation datasets were next subjected to the QN.BMIQ normalization procedure [149]. More specifically, we first applied quantile normalization as implemented in the lumi R package to remove systematic effects between samples. Next, we applied the β-mixture quantile normalization (BMIQ) procedure [150] as implemented in the wateRmelon R package [151] to normalize beta values of type 1 and type 2 design probes within the Illumina arrays.

Next, we removed confounding effects in DNA methylation data by fitting the model *median methylation M value ∼ neuron.proportions + batch + sample.plate array + ageAtDeath + sex* and extracting residuals from this model, which are the methylation residuals. Similarly, we also removed potential confounding effects in RNA-seq data by fitting model *log_2_ (normalized FPKM values + 1) ∼ ageAtDeath + sex + markers for cell types*. The last term *markers for cell types* included multiple covariate variables to adjust for the multiple types of cells in the brain samples. More specifically, we estimated expression levels of genes that are specific for the main five cell types present in the CNS: ENO2 for neurons, GFAP for astrocytes, CD68 for microglia, OLIG2 for oligodendrocytes, and CD34 for endothelial cells, and included these as variables in the above linear regression model, as was done in a previous large study of AD samples [6]. The residuals extracted from this model are the gene expression residuals.

TF activities were calculated using the collection of brain-specific TF Regulons (downloaded from ChEA3 software website at https://maayanlab.cloud/chea3/assets/tflibs/brain.TFs.gmt) and gene expression residuals, using the R package GSVA (Gene Set Variation Analysis) with the following parameters: method = “gsva”, kcdf = “Gaussian”, abs.ranking = TRUE, min.sz = 5, max.sz = Inf, mx.diff = TRUE, ssgsea.norm = TRUE.

#### Creating triplet datasets

For supervised MethReg analysis, we considered the 3,751 differently methylated CpGs associated with Braak stage identified in Zhang et al. (2020) [89]. First, these CpGs were filtered to remove probes with beta-values IQR < 0.03, resulting in 2,761 CpGs. For CpGs in the promoter regions, the function create_triplet_distance_based with parameters target.method = “genes.promoter.overlap” was used to link each of the 1002 CpGs in promoter regions (within ± 2 kb regions around the TSS) to the target gene with promoter region that overlaps with the CpG. In distal analysis, the function create_triplet_distance_based with parameters target.method = “nearby.genes” and target.num.flanking.genes = 5 were used to link each of the 1,759 CpGs in distal regions (beyond ± 2 kb regions around TSS of the target genes) to five genes upstream and downstream of the CpG. Alternatively, in regulon-based analysis, the function create_triplet_regulon_based was used to link CpGs to the TF-target gene pairs in the brain-specific regulons (downloaded from https://maayanlab.cloud/chea3/). In all three analyses, we obtained TF binding sites information from the REMAP2020 database and linked CpGs to TFs with binding sites within 250 bp. The CpG-TF pairs were then combined with CpG-target gene pairs (promoter and distal analyses) or TF-target pairs (regulon-based analysis) to obtain CpG-TF-target triplets. In all analyses, we specified the parameter max.distance.region.target = 10^6 to obtain triplets for which the CpG is within 1M bp distance from the target gene. These analyses were performed using hg19 genome assembly coordinates.

#### Overlap with cis-regulatory elements in enhancer database

We assessed the overlap between regions in significant triplets (CpG location +/- 250bp) with enhancer regions in EnhancerAtlas 2.0 database [152]. The following brain/neuron specific tissues or cell lines for Homo sapiens (hg19) were considered: Astrocyte, hNCC, KELLY, BE2C, ESC_NPC, NGP, SK.N.MC, Cerebellum, Gliobla, SH.SY5Y, SK.N.SH_RA, T98G, Fetal_brain, U87, H54, SK.N.SH, SK.N.MC, NH.A, Macrophage, ESC_neuron.

## Acknowledgement

This research was supported by US National Institutes of Health grants R21AG060459 (L.W), R01AG061127 (L.W.), and R01AG062634 (E.R.M, L.W.). The ROSMAP study data were provided by the Rush Alzheimer’s Disease Center, Rush University Medical Center, Chicago. Data collection was supported through funding by NIA grants P30AG10161, R01AG15819, R01AG17917, R01AG30146, R01AG36836, U01AG32984, U01AG46152, the Illinois Department of Public Health, and the Translational Genomics Research Institute.

## Availability of data and materials

The MethReg R package is available from the Bioconductor repository at https://bioconductor.org/packages/MethReg/.

## Author Contributions

L.W., T.C.S., J.Y. designed the computational analysis. T.C.S., L.W. analyzed the data. L.W., J.Y., E.R.M, X.C. contributed to interpretation of the results. T.C.S, L.W. wrote the paper, and all authors participated in the review and revision of the manuscript. L.W. conceived the original idea and supervised the project.

## Ethics approval and consent to participate

Not applicable.

---

**Supplementary Figure 1.**
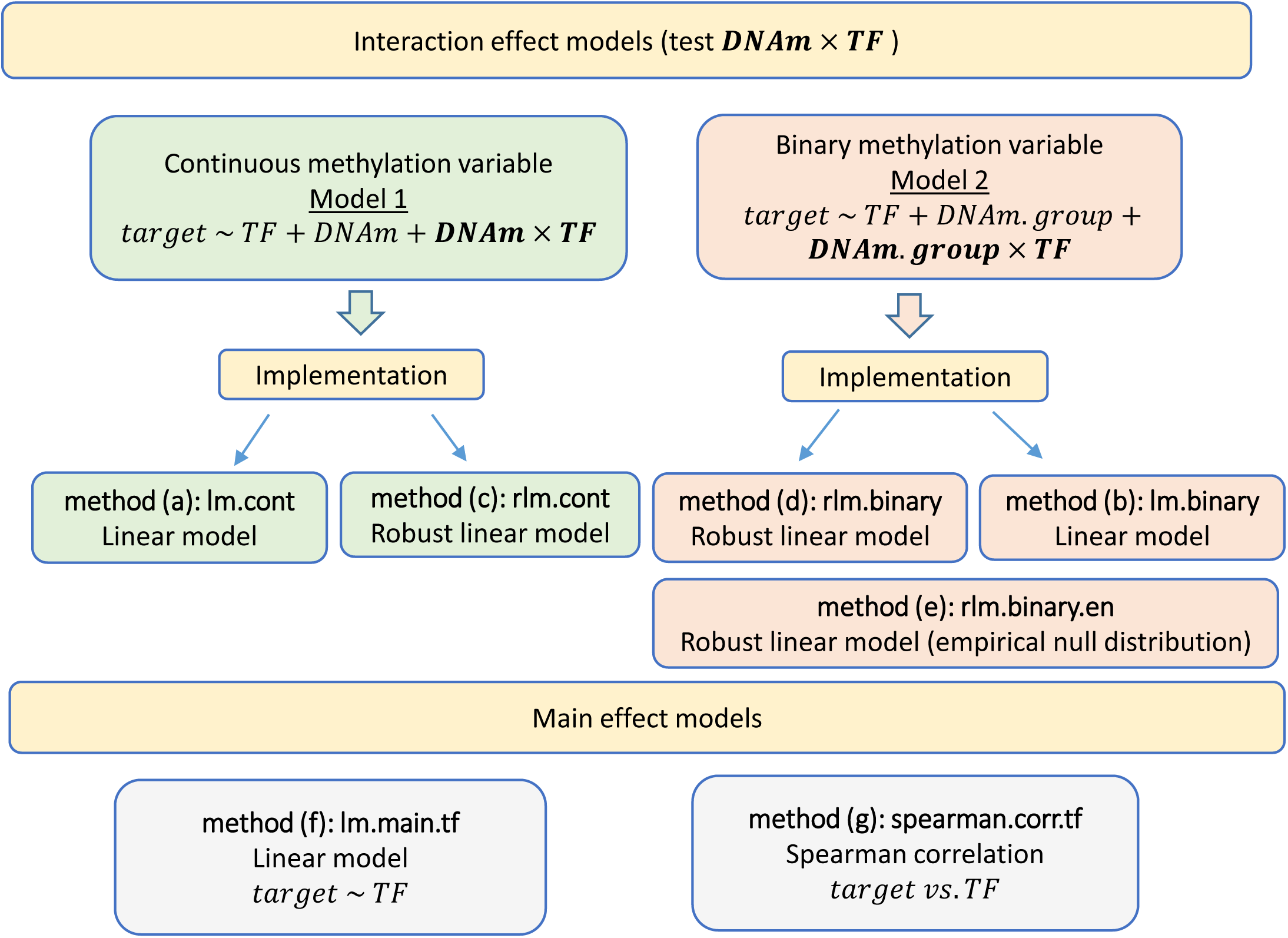
Methods compared in simulation study.

**Supplementary Figure 2.**
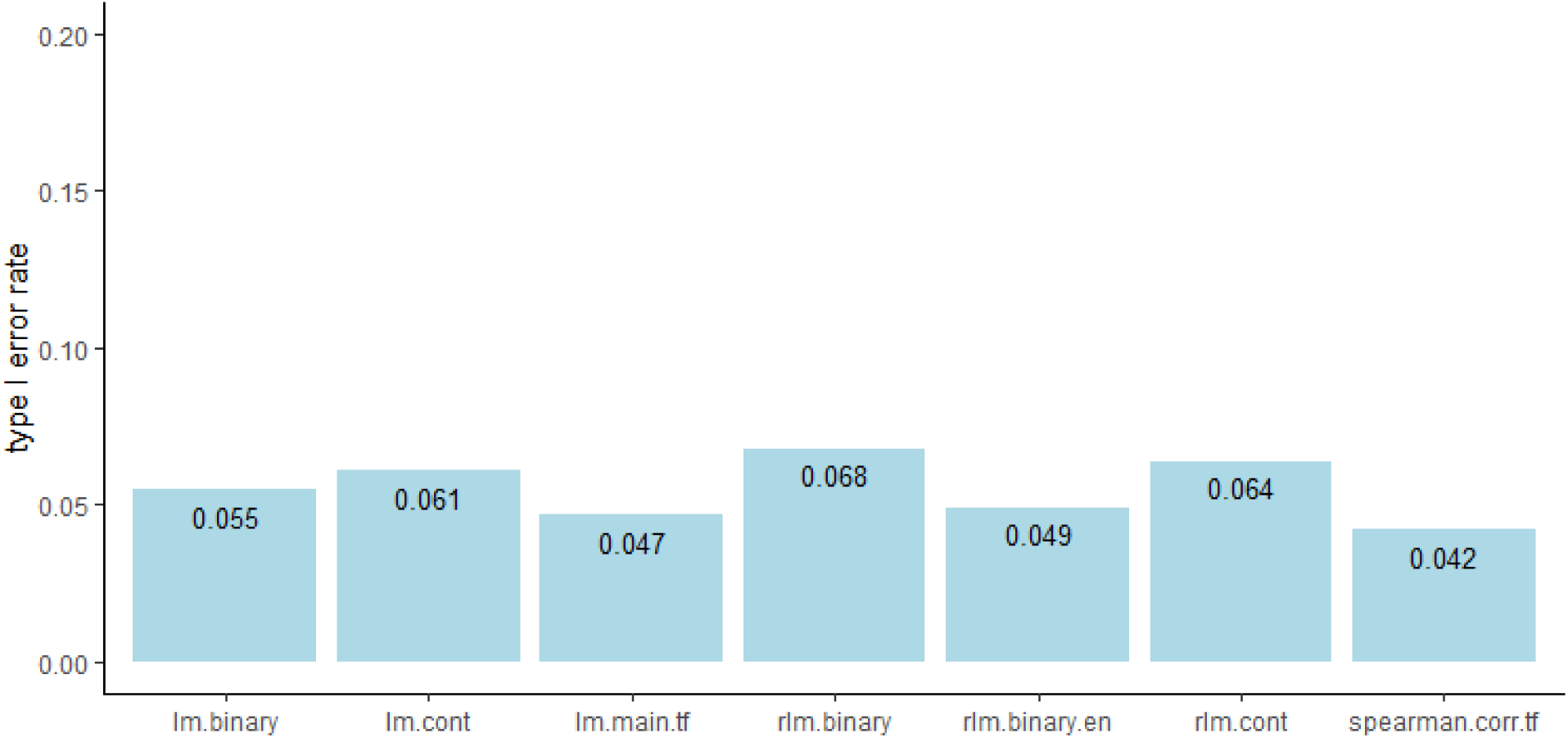
Type I error rates of different methods in simulated datasets. lm.binary = linear model implementation of Model 2; lm.cont = linear model implementation of Model 1; lm.main.tf = linear model *gene expression ∼ TF*; rlm.binary = robust linear model implementation of Model 2; rlm.binary.en = same as rlm.binary model, except P-values were estimated using empirical null distribution; rlm.cont = robust linear model implementation of Model 1; spearman.corr.tf = Spearman correlation between gene expression and TF expression; Model 1: *gene expression ∼ TF + DNAm + TF*DNAm*; Model 2: *gene expression ∼ TF + DNAm.group + TF*DNAm.group*

**Supplementary Figure 3.**
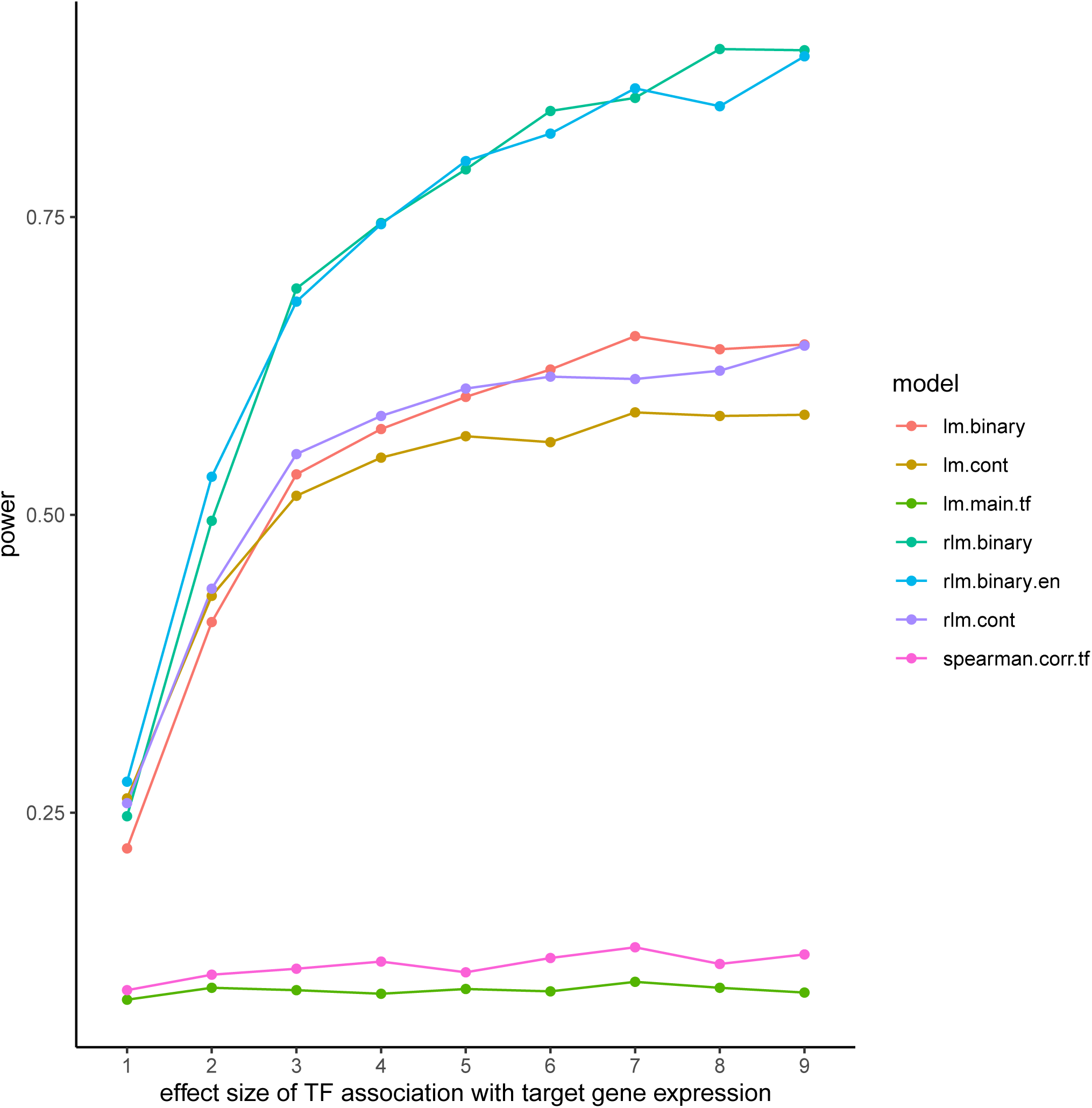
Power of different methods in simulation study. lm.binary = linear model implementation of Model 2; lm.cont = linear model implementation of Model 1; lm.main.tf = linear model *gene expression ∼ TF*; rlm.binary = robust linear model implementation of Model 2; rlm.binary.en = same as rlm.binary model, except P-values were estimated using empirical null distribution; rlm.cont = robust linear model implementation of Model 1; spearman.corr.tf = Spearman correlation between gene expression and TF expression; Model 1: *gene expression ∼ TF + DNAm + TF*DNAm;* Model 2: *gene expression ∼ TF + DNAm.group + TF*DNAm.group*

**Supplementary Figure 4.**
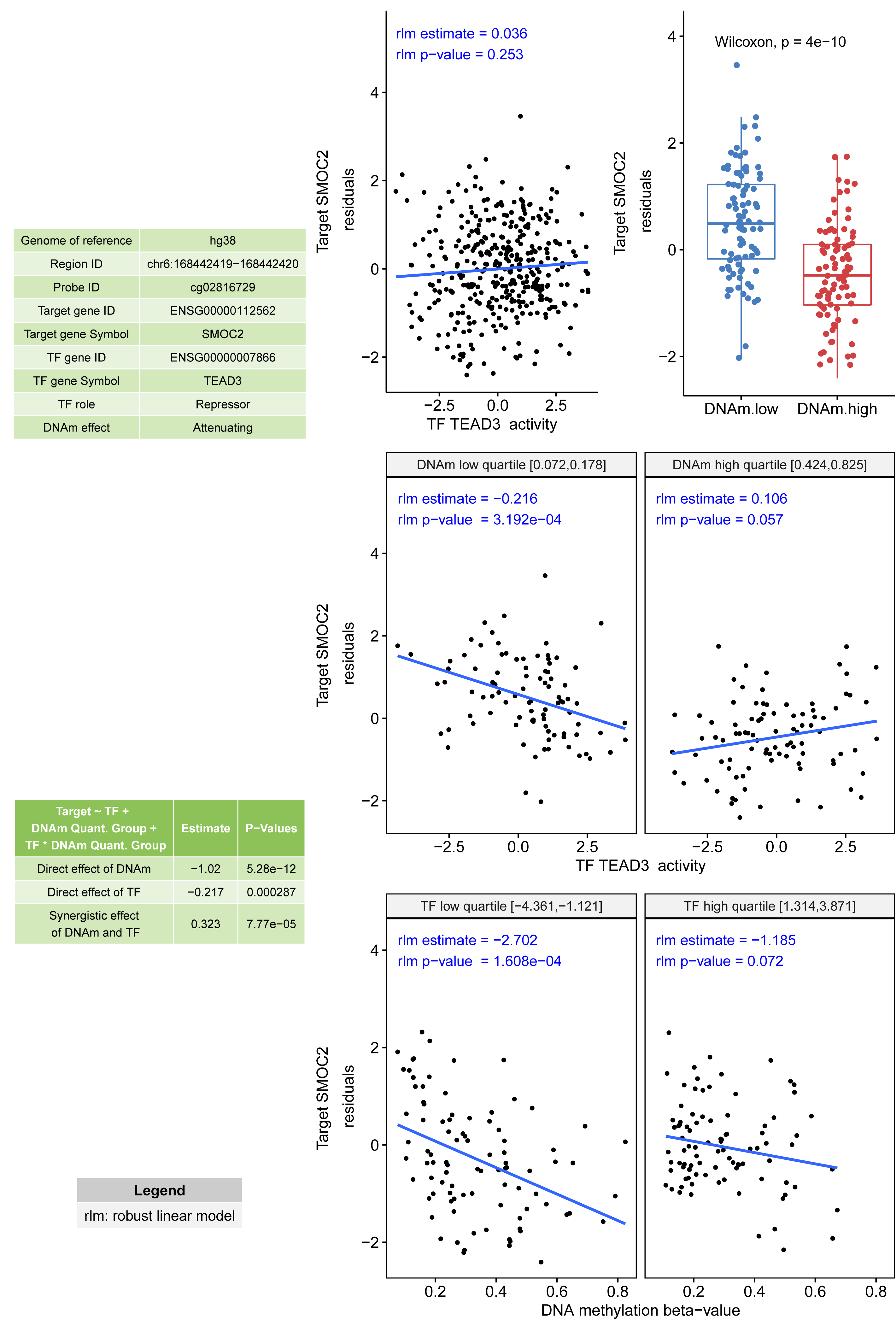
An example of CpG methylation attenuating TF activities, from unsupervised MethReg analysis of TCGA COAD-READ samples.

**Supplementary Figure 5.**
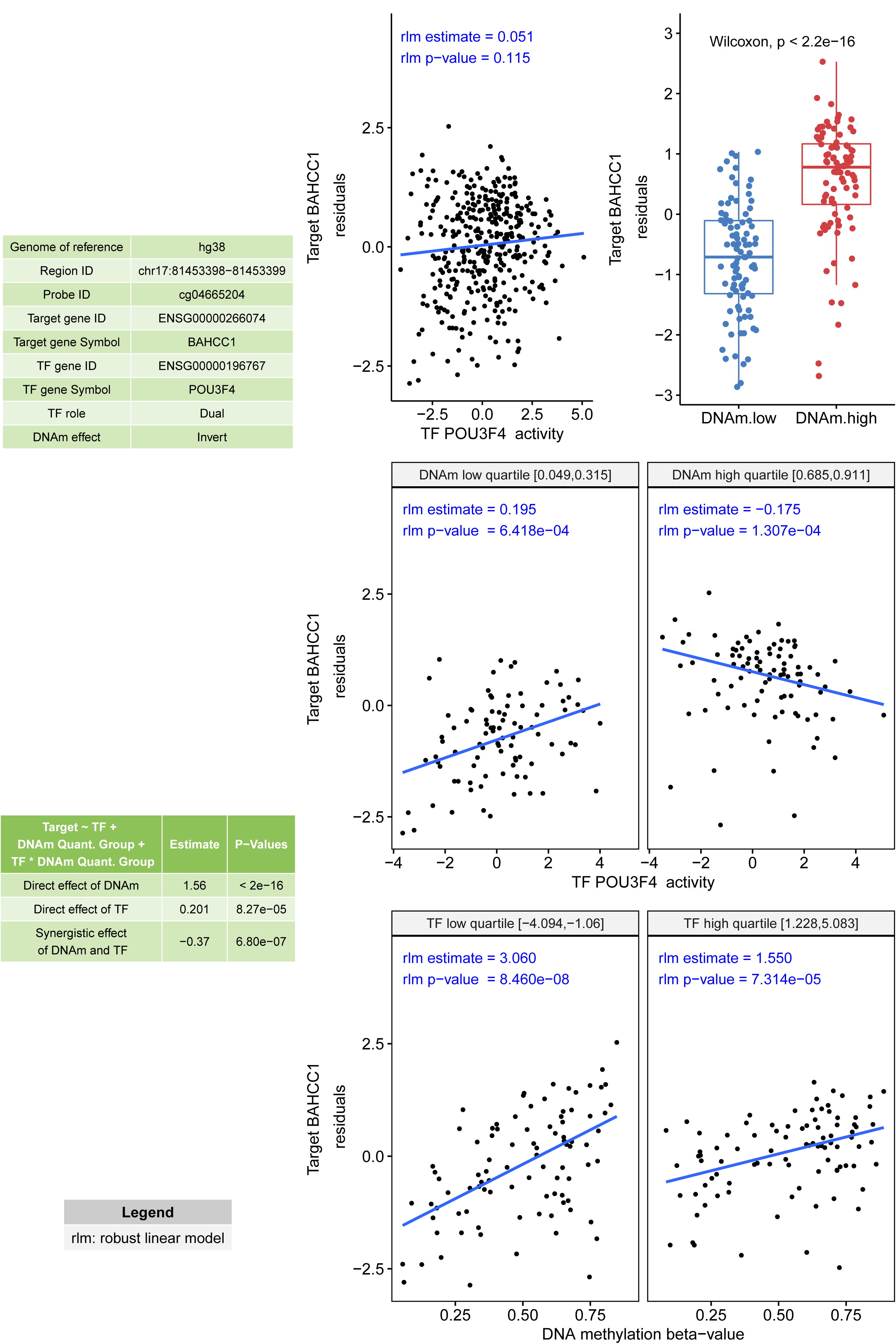
An example of TF with dual modes of regulations depending on CpG methylation levels, from unsupervised MethReg analysis of TCGA COAD-READ datasets.

**Supplementary Figure 6.**
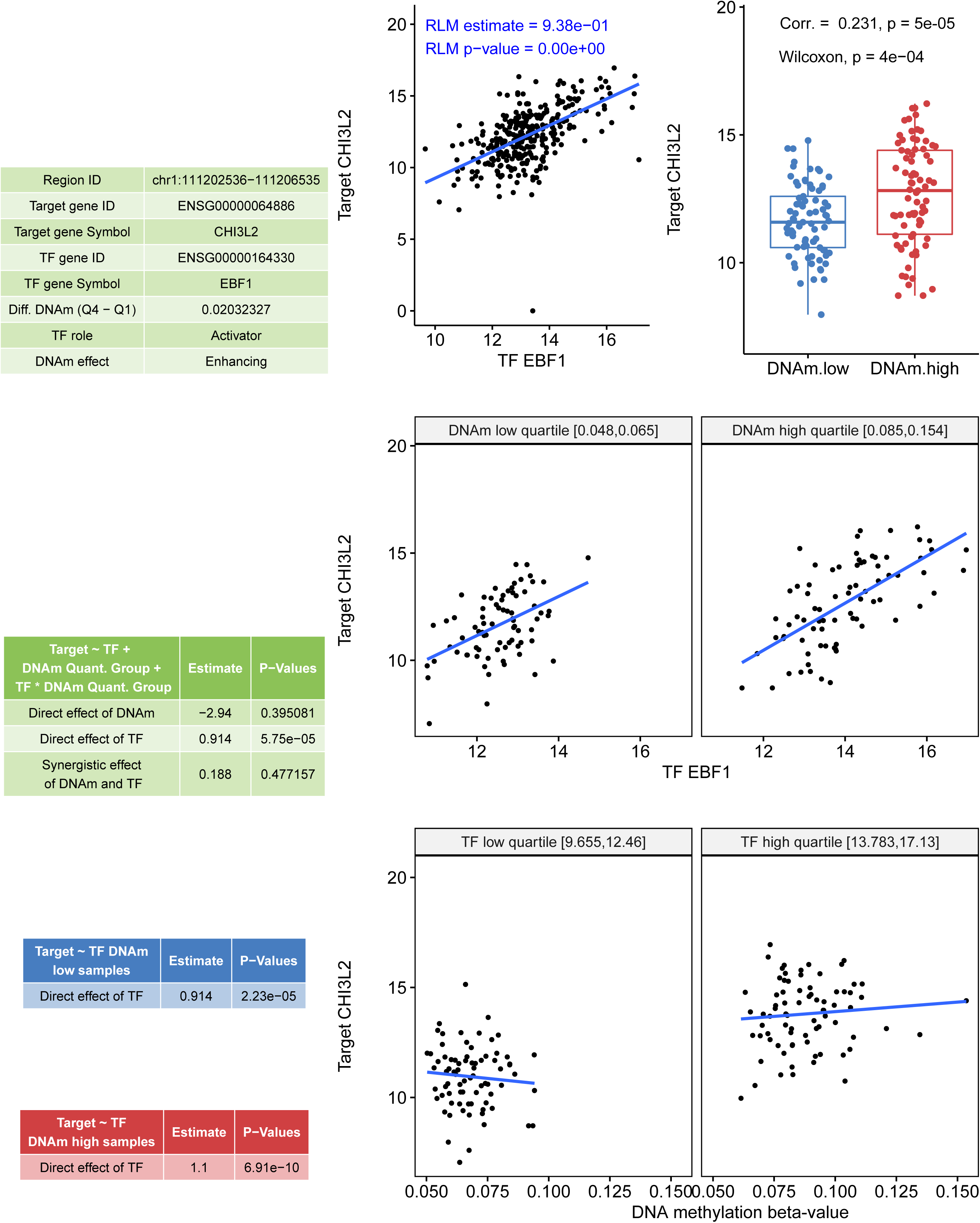
Example of confounding TF effect. The target gene expression is mainly driven by the TF EBF1, and not by DNA methylation, even though a highly significant methylation-target gene association was observed.

**Supplementary Figure 7.**
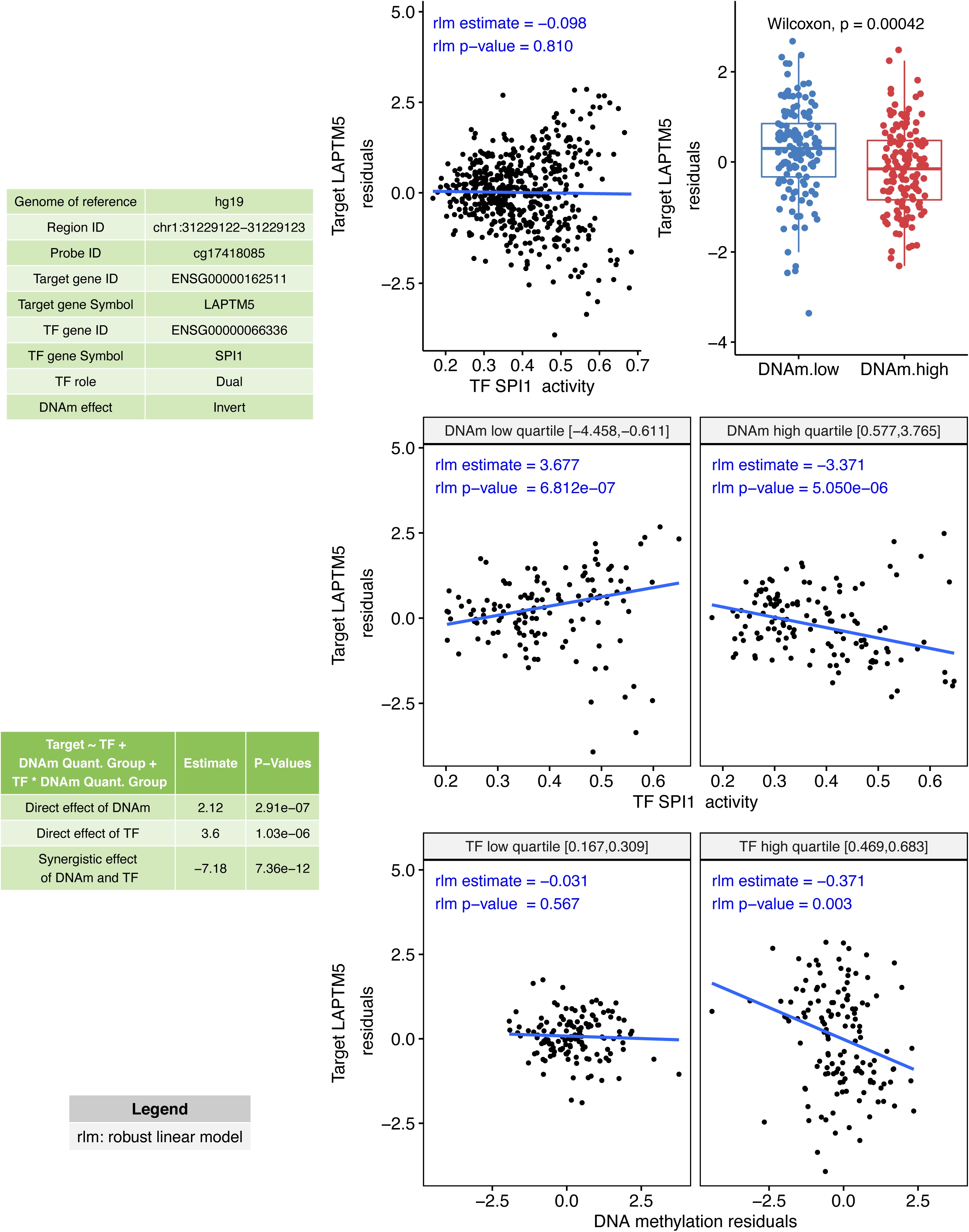
An example of TF having dual regulatory roles depending on CpG methylation levels, from supervised MethReg analysis ROSMAP dataset.

**Supplementary Figure 8.**
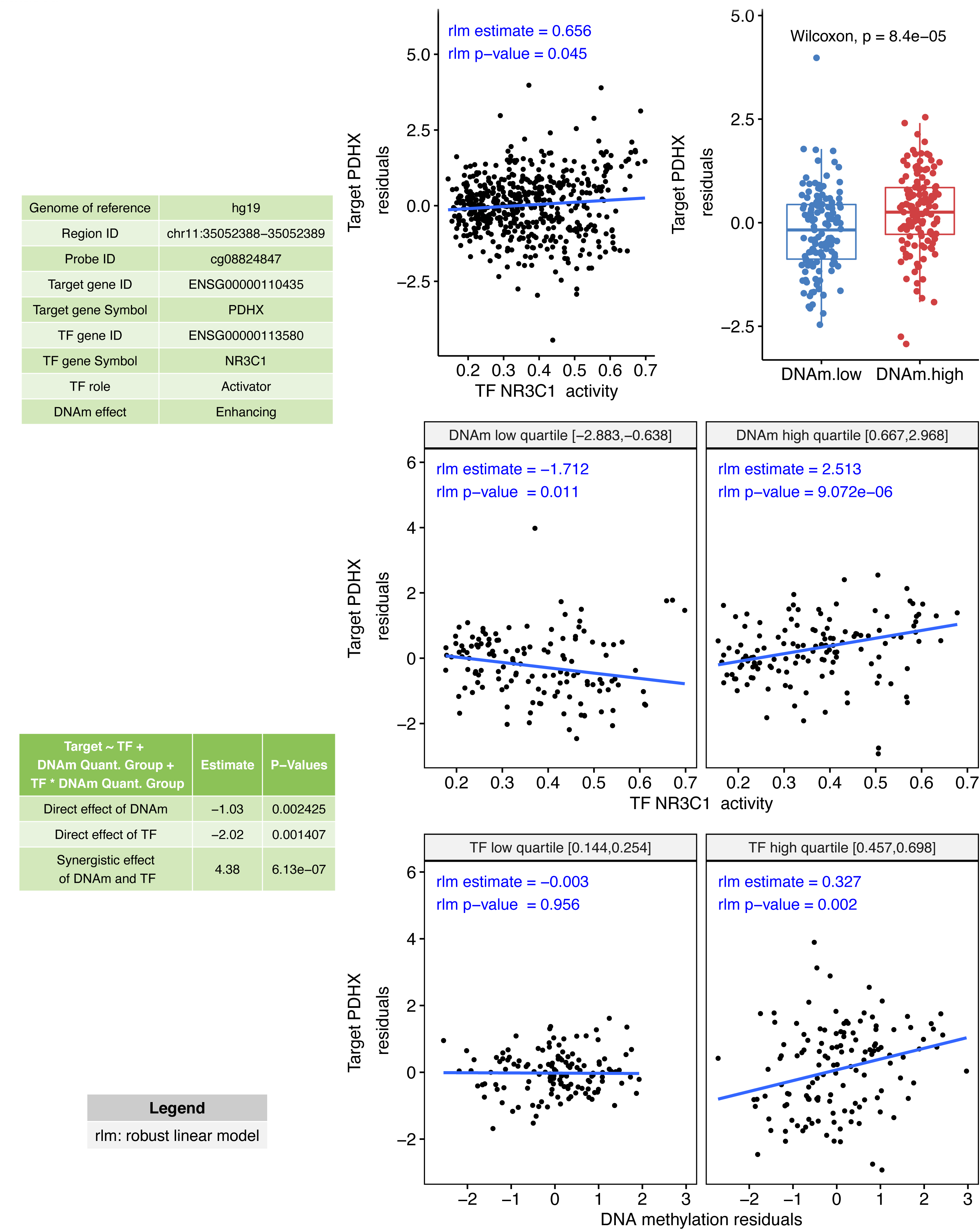
An example of CpG methylation enhancing TF activities, from supervised MethReg analysis of ROSMAP dataset.

